# FDA drug repurposing uncovers modulators of dopamine D_2_ receptor localization via disruption of the NCS-1 interaction

**DOI:** 10.1101/2025.05.29.656787

**Authors:** D Muñoz-Reyes, L Aguado, S Arroyo-Urea, C Requena, S Pérez-Suárez, S Sánchez-Yepes, J Argerich, C Miró-Rodríguez, E Ulzurrun, E Rodríguez, J García-Nafría, NE Campillo, A Mansilla, MJ Sánchez-Barrena

**Author notes:** Author contributions:* DMR, SPS and CMR carried out protein expression, purification, *in vitro* biophysical experiments and crystallization; SPS generated the constructs for *in vitro* assays, PLA and localization studies; and DMR collected diffraction data, performed model building, refinement and structural analysis. All this work was supervised by MJSB. LA, SSY, ER performed PLA assays, cellular localization and trafficking studies under the direction of AM. CR and EU, supervised by NEC, performed virtual screening studies and molecular dynamics simulations. SAU and JA carried out functional BRET2 and NanoBiT assays, with supervision of JGN. MJSB, AM, NEC and JGN designed the research and discussed the manuscript. MJSB contributed with the initial conception, coordinated with AM and NEC. MJSB and AM wrote the manuscript and included contributions from all authors. The authors declare no conflict of interest. This article contains supporting information. Data deposition:* The atomic coordinates and structure factors of the NCS-1 in complex with the FDA ligands have been deposited in the Protein Data Bank, https://www.pdb.org/, with PDB codes: NCS-1/AZS (9GTO), NCS-1/ATV (9GU6), NCS-1/VLZ (9GU8).

## Abstract

Dopamine D_2_ receptor (D_2_R) regulates key aspects of motor control, cognition, and reward. Its function depends not only on ligand binding and signaling efficacy, but also on the dynamic control of receptor localization at the cell surface. Neuronal Calcium Sensor 1 (NCS-1) is a calcium binding protein which directly interacts with D_2_R in a Ca^2+^-dependent manner. Here, we investigated the regulatory role of NCS-1 in D_2_R localization and function. We found that NCS-1 promotes the trafficking of D_2_R to the plasma membrane through a mechanism dependent on active exocytosis. Functional signaling assays confirmed that NCS-1 does not alter the canonical receptor pharmacology. Using a library of FDA-approved drugs, a structure-based drug-repurposing strategy was designed to find protein-protein interaction modulators that allowed the exploration of the NCS-1/D_2_R interface as a new pharmacological target. Azilsartan medoxomil, atorvastatin, and vilazodone disrupted its interaction with D_2_R, reducing receptor surface expression in cells. Crystallography and molecular dynamics simulations revealed their mechanism of action. These compounds target the NCS-1 hydrophobic crevice and overlap the D_2_R binding site, perturbing the dynamics of the regulatory helix H10 in NCS-1. These findings uncover a previously unexploited intracellular mechanism for modulating D_2_R function and highlight the potential of targeting protein–protein interactions for therapeutic purposes. Our results provide a framework for fine-tuning dopaminergic tone through receptor localization mechanisms, offering an alternative strategy to conventional approaches based on receptor blockade or direct agonism.

**SIGNIFICANCE:** FDA-approved drugs targeting the calcium sensor NCS-1 selectively disrupt its interaction with the dopamine D_2_ receptor, offering a new strategy to modulate receptor trafficking without altering receptor signaling.

## INTRODUCTION

The dopamine D_2_ receptor (D_2_R) is a member of the G protein–coupled receptor (GPCR) family and transduces signals primarily through the coupling and activation of heterotrimeric Gαβγ proteins (G_i/O_ type) and β-arrestins (1, 2). D_2_R is a key pharmacological target in the treatment of various neuropsychiatric disorders, including Tourette’s syndrome, schizophrenia, Parkinsońs disease and bipolar disorders, conditions commonly associated with imbalance in dopaminergic transmission. This is underscored by the fact that D_2_-like receptors are the major target for drugs in the treatment of Parkinsońs disease (agonists) and the majority of antipsychotic and neuroleptic drugs (antagonists and partial agonists), with D_2_R representing the predominant isoform expressed in the brain. In addition, human genetic variants in the D_2_R have been shown to yield motor, cognitive, and neuropsychiatric deficits (3–10).

The density of postsynaptic dopamine D_2_ receptors critically determines neuronal sensitivity to dopamine, thereby influencing both the intensity and duration of dopaminergic signaling. Proper neural communication relies on the maintenance of an optimal level of D_2_R surface expression: insufficient receptor availability may attenuate dopamine-mediated responses, whereas excessive expression can result in heightened sensitivity and dysregulated signaling activity. In addition to their postsynaptic role, D_2_Rs also function as presynaptic autoreceptors located on dopaminergic neurons. Activation of these autoreceptors initiates intracellular signaling pathways that inhibit further dopamine release, establishing a negative feedback loop essential for maintaining dopaminergic homeostasis and preventing neurotransmitter overaccumulation (11, 12).

The functional availability of D_2_Rs at both pre- and postsynaptic sites is tightly regulated by receptor trafficking processes, including insertion into the membrane, internalization, and recycling. These dynamic mechanisms control the number of receptors accessible at the cell surface and are therefore central to dopaminergic regulation (13). Among the proteins implicated in modulating D_2_R trafficking is Neuronal Calcium Sensor 1 (NCS-1), which has been shown to influence receptor localization, although the precise molecular mechanisms remain incompletely defined (14). Elucidating these regulatory pathways is critical for understanding dopamine-related neural function and its dysregulation in neuropsychiatric disorders.

NCS-1 is a high-affinity calcium binding protein that plays diverse roles in the nervous system, including synaptic transmission and plasticity, due to its ability to interact and regulate multiple protein targets. NCS-1 regulates the activity of Ca^2+^ channels such as the inositol 1,4,5-trisphosphate receptors (InsP_3_Rs) and several proteins implicated in G-protein signaling: dopamine D_2_ receptor, GRK2 kinase or the Gα chaperone and GEF Ric-8A, through both in a Ca^2+^-dependent and -independent mechanisms (14–16). Dysregulation of NCS-1 has been associated with several neuropathological processes including X-linked intellectual disability, autism, schizophrenia, and bipolar disorder (17–20). Notably, elevated NCS-1 expression has been reported in the prefrontal cortex of individuals with schizophrenia and bipolar disorder (18). Similarly, an increase in post-synaptic D_2_R levels has been observed in antipsychotic-free schizophrenia patients. Collectively, these findings highlight the importance of investigating the regulatory role of NCS-1 in D_2_R function and its impact on cellular localization, receptor signaling and pharmacology.

Previous studies have identified NCS-1 as a potential therapeutic target, where modulation of the NCS-1/Ric-8A interaction by small molecules can restore synaptic function, showing promise in models of fragile X syndrome and Alzheimeŕs disease (21–23). These compounds exert their effects by modulating the conformational dynamics of the C-terminal helix H10 of NCS-1, which serves as a regulatory gatekeeper for Ric-8A binding, which inserts a long helix into an elongated and hydrophobic cavity under resting intracellular Ca^2+^ state (16, 21). In contrast, NCS-1 interacts with the dopamine D_2_ receptor under elevated intracellular Ca^2+^ concentrations. In this case, the helix H10 inserts into the elongated crevice, reshaping NCS-1 molecular surface to create the D_2_R recognition interface, where two intracellular H8 helices bind to (Figure 1) (14, 24, 25).

**Figure 1:**
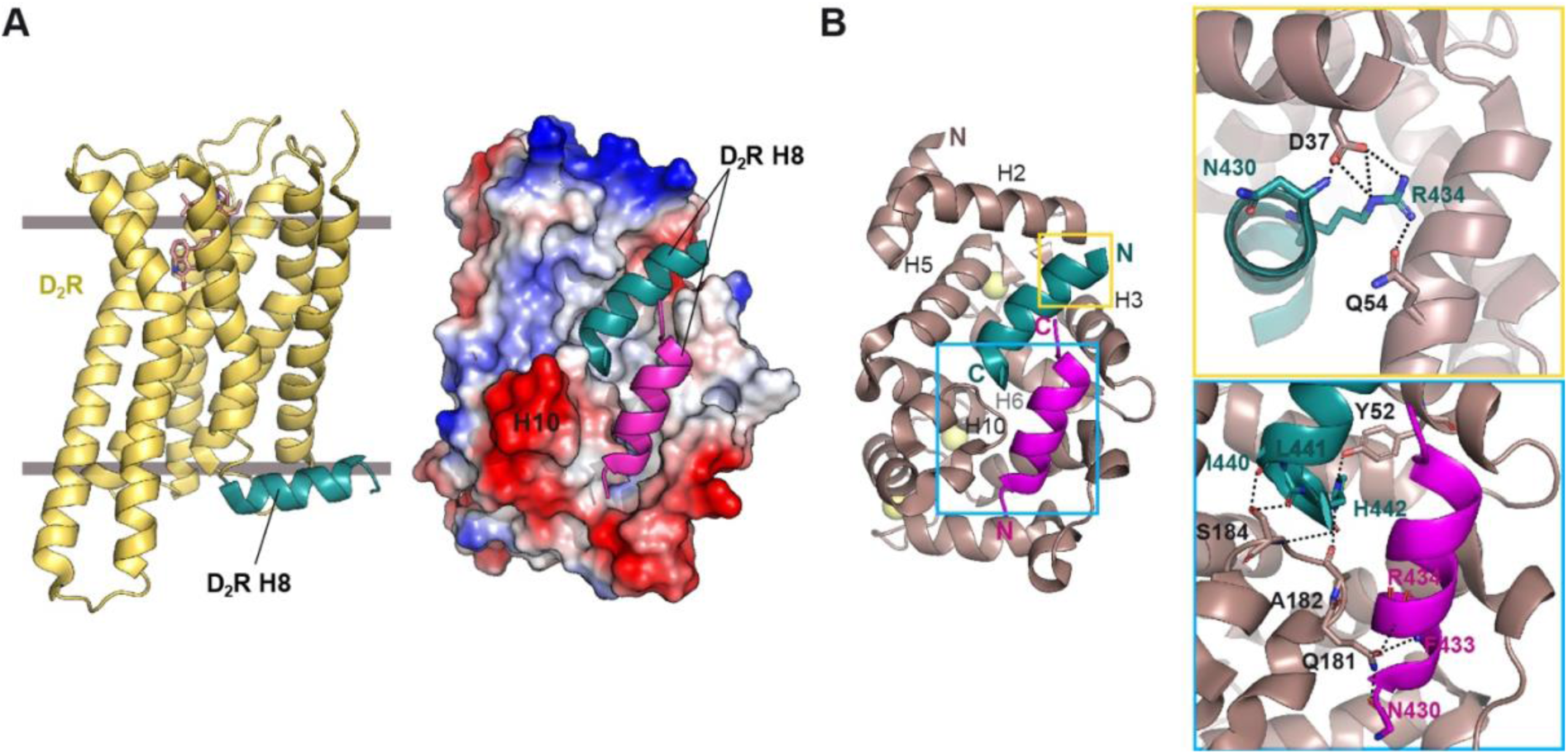
NCS-1 helix H10 participates in dopamine D_2_ receptor recognition. **(A)** Left: The structure of D_2_R bound to an agonist (PDB: 6VMS, (74)) and showing in green the intracellular helix H8. Right: Molecular surface representation of NCS-1 bound to D_2_R H8 helices (PDB: 5AER, (25)). **(B)** Details of the strong polar interactions occurring at the ends of the H8 helices. Sidechains not involved in H-bond interactions with NCS-1 are not shown.

Building on these structural insights, we implemented a structure-based drug repurposing approach using an FDA-approved compound library to identify small-molecule modulators of the NCS-1/D_2_R interaction. These compounds serve not only as molecular probes to investigate the role of NCS-1 in D_2_R trafficking and signaling, but also as potential therapeutic hits for the treatment of neuropsychiatric disorders such as schizophrenia and bipolar disorder.

## RESULTS

### NCS-1 regulates the subcellular localization of D_2_R

To assess the effects of NCS-1 on D_2_R localization, cells were transfected, and sequential immunostaining was performed to detect membrane-bound D_2_R in red and cytoplasmic D_2_R in green. Quantification of subcellular localization based on the confocal images showed that under basal conditions, D_2_R is mainly localized in the cytoplasm, whereas NCS-1 overexpression promotes a significant accumulation of D_2_R at the plasma membrane (Figure 2A,C).

**Figure 2:**
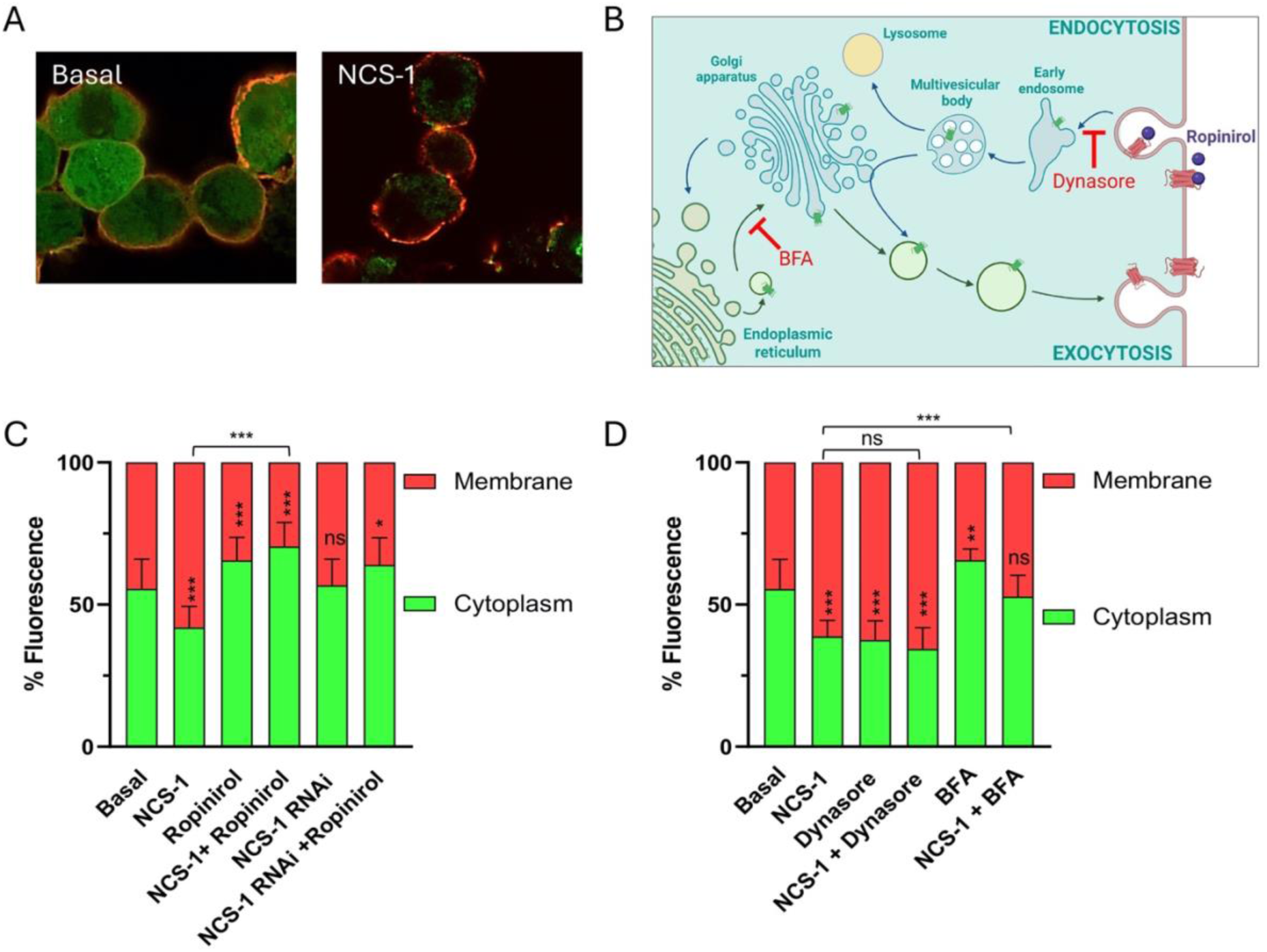
Role of NCS-1 in the subcellular localization of D_2_R. **(A)** Representative confocal images of HEK293 cells transfected with D_2_R alone (Basal) or co-transfected with D_2_R and NCS-1 (NCS-1). D_2_R is FLAG-tagged, and a sequential immunofluorescence protocol was used to differentially label plasma membrane-localized D_2_R (red) and intracellular D_2_R (green). **(B)** Schematic diagram illustrating D_2_R trafficking via endocytic and exocytic pathways, along with the targets of the pharmacological agents used in the study: Ropinirole (a D_2_R agonist that promotes receptor internalization), Dynasore (an endocytosis inhibitor), and Brefeldin A (BFA, an exocytosis inhibitor). Illustration created with BioRender.com. **(C)** Quantification of D_2_R fluorescence distribution between membrane (red) and cytoplasm (green) in HEK293 cells transfected with D_2_R alone or with NCS-1 or NCS-1 RNAi. Ropinirol is used as a D_2_R agonist. **(D)** Effect of Dynasore and BFA on NCS-1-mediated D_2_R redistribution. Data are expressed as the percentage of total D_2_R fluorescence localized at the membrane (red) and in the cytoplasm (green). Asterisks within bars indicate statistical comparison with the Basal condition (D_2_R alone), while horizontal bars above the graphs indicate comparisons between specific experimental conditions. Statistical significance: ns = not significant, *p < 0.05, **p < 0.01, ***p < 0.001.

Ropinirole, a D_2_R agonist known to trigger internalization, reversed this effect when applied to NCS-1-overexpressing cells, suggesting that NCS-1 promotes D_2_R membrane stabilization, which can be counteracted by agonist-induced endocytosis. Interestingly, knockdown of HEK293 endogenous NCS-1 expression via RNA interference led to a shift of D_2_R back to the cytoplasm, further supporting the role of NCS-1 in plasma membrane localization (Figure 2B,C).

To dissect the underlying trafficking mechanisms, we used Dynasore (endocytosis inhibitor) and Brefeldin A (BFA, exocytosis inhibitor) in combination with NCS-1 overexpression (Figure 2B,D) (26). Dynasore did not significantly affect NCS-1-induced membrane localization of D_2_R, suggesting that NCS-1 does not act primarily by blocking endocytosis. In contrast, BFA treatment prevented the NCS-1-induced increase in membrane D_2_R, indicating that NCS-1 enhances D_2_R membrane localization through a BFA-sensitive exocytic pathway.

Overall, these findings indicate that NCS-1 promotes the trafficking of D_2_R to the plasma membrane through a mechanism dependent on active exocytosis rather than inhibition of endocytosis. This highlights NCS-1 as a positive regulator of D_2_R surface expression and may have implications for dopaminergic signaling regulation.

### The role of NCS-1 in regulating dopamine D_2_ receptor function

The dopamine D_2_ receptor belongs to the GPCR family and hence signals by coupling to and activating heterotrimeric Gαβγ proteins and β-arrestins (2). It is currently unclear whether binding of NCS-1 at the intracellular helix H8 affects the receptor pharmacological properties, since its binding potentially poses a steric conflict with intracellular transducers (Figure 1). For this purpose, we performed signaling assays in HEK293T cells, titrating quinpirole, a widely used D_2_R agonist, to obtain potency (pEC_50_) and efficacy (E_max_) pharmacological values in three different assays that included Gαβγ activation (using BRET2 assays) (27), miniGα recruitment (using Nanobit assays) (28) and β-arrestin recruitment to the receptor (using split Nanoluc assays) (29), all in the presence and absence of co-transfected NCS-1. Overall, the potency of quinpirole in the three assays was virtually identical in the absence and presence of NCS-1 (Figure 3 and Supplementary Figure 1), highlighting that NCS-1 does not have an impact in the canonical functions of the receptor. Additionally, efficacy was also identical when monitoring Gαβγ activation and the recruitment of β-arrestin (Figure 3), hence also independent of the presence of NCS-1. These signaling assays monitor signaling processes immediately at the receptor, lacking signal amplification and thus are not sensitive to minor changes in receptor expression (27). Therefore, we do not expect to see a difference in efficacy due to the increase in surface expression of the D_2_R caused by NCS-1 co-transfection. An increase in efficacy was seen in the Nanobit assay upon NCS-1 co-transfection (Supplementary Figure 1), however it seems a much larger efficacy effect than what would be expected from the increase in surface expression seen with confocal microscopy (Figure 2). Although we cannot fully explain the increase in efficacy on these assays, miniGα proteins have been shown to disrupt receptor endocytic trafficking (30), which could, together with NCS-1, yield complex trafficking effects, only observed when using such protein chimeras but not when using active (non-dominant negative) Gαβγ proteins and β-arrestins. Overall, we conclude that over-expressing NCS-1 does not have an impact on the pharmacological profile of the D_2_R in this cellular model system.

**Figure 3:**
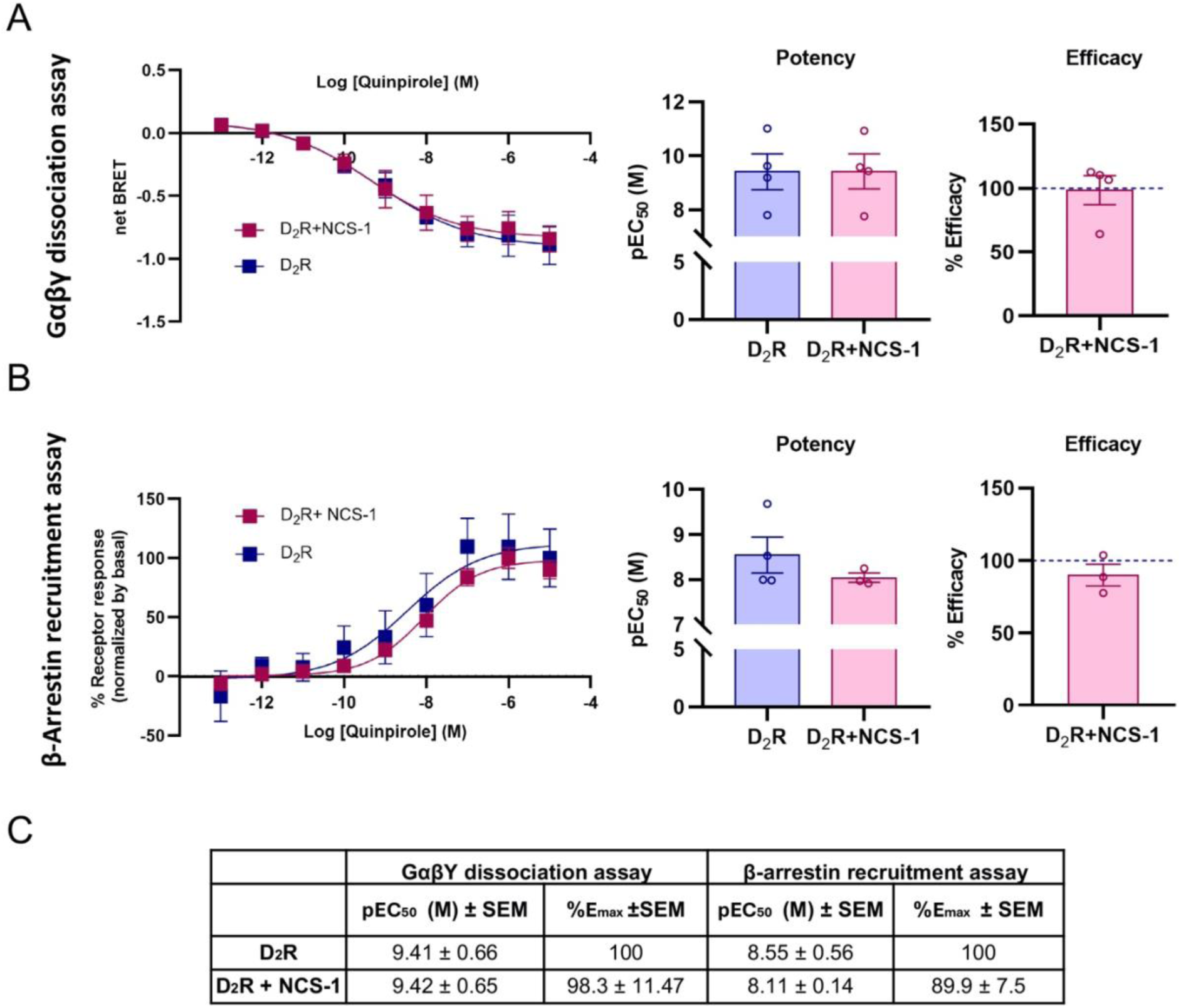
Effect of NCS-1 on D_2_ receptor-mediated G protein activation and β-arrestin recruitment. **(A)** Concentration-response curves and bar plots of D_2_R (blue) and D_2_R co-transfected with NCS-1 (pink) upon G_OA_ activation with quinpirole using cellular BRET2 assays. **(B)** Concentration-response curves and bar plots of D_2_R (blue) and D_2_R co-transfected with NCS-1 (pink) upon β-Arrestin recruitment with quinpirole. **(C)** Summary table with pEC_50_ and E_max_ values derived from the Gαβγ dissociation and β-arrestin recruitment assays. Data are presented as means ± SEM and pEC_50_ and E_max_ values of all data are derived from four independent experiments performed in technical triplicates except for D_2_R + NCS-1 in β-arrestin recruitment assay (n=3). E_max_ bar plots are displayed as normalized efficacy to D_2_R (dashed blue line). Source data are provided as a Source Data File.

### Virtual screening to find FDA compounds targeting the NCS-1/D_2_R interface

To identify compounds capable of modulating the NCS-1/D_2_R protein-protein interaction (PPI) interface, virtual screenings were conducted using the FDA-approved drug library comprising 2143 compounds. Given the structural characteristics of NCS-1, particularly the key role of helix 10 (H10) in mediating interactions with other proteins (21, 31, 22), we proposed conducting the virtual screening using different structural models of NCS-1. To this end, a structural analysis was performed to select the most suitable NCS-1 template models for screenings.

NCS-1 contains a dynamic C-terminal helix (H10) that shapes the exposed hydrophobic crevice, the primary binding site for target proteins (25, 21, 15, 16). The crystal structure of NCS-1 in complex with the cytosolic C-terminal H8 helix of D_2_R (PDB 5AER; (25)) revealed that H10 defines two cavities: an upper cavity where a D_2_R H8 is deeply inserted and a lower cavity where a second copy of the D_2_R H8 is found. In addition, NCS-1 H10 directly interacts with D_2_R H8, with residues D37, Y52 and Q181 establishing strong interactions and thus, playing key roles in receptor recognition (Figure 1). Given the importance of NCS-1 H10 in D_2_R binding, we selected two NCS-1 structural templates for virtual screenings: one with H10 inserted in the crevice (PDB: 5AAN; (21)) and another with H10 exposed (PDB: 6QI4, (22)). Furthermore, a filtering criterion was applied to prioritize molecules interacting with D37, Y52 and Q181.

A hierarchical virtual screening approach was employed, targeting the two proposed structural models of NCS-1. Both targets were subjected to a three-staged virtual screening protocol consisting of a preliminary docking study by using the SP Glide docking algorithm, a second docking by applying the XP Glide docking algorithms and a final rescoring by applying the Prime MM-GBSA method. For each screened target, at least the 20 best ranked FDA drugs according to the MM-GBSA score were preliminarily selected. Finally, the compounds were filtered based on their interactions with key residues (D37, Y52 and Q181) in order to identify molecules with the most favorable interactions (Figure 4 and Supplementary Table 1).

**Figure 4:**
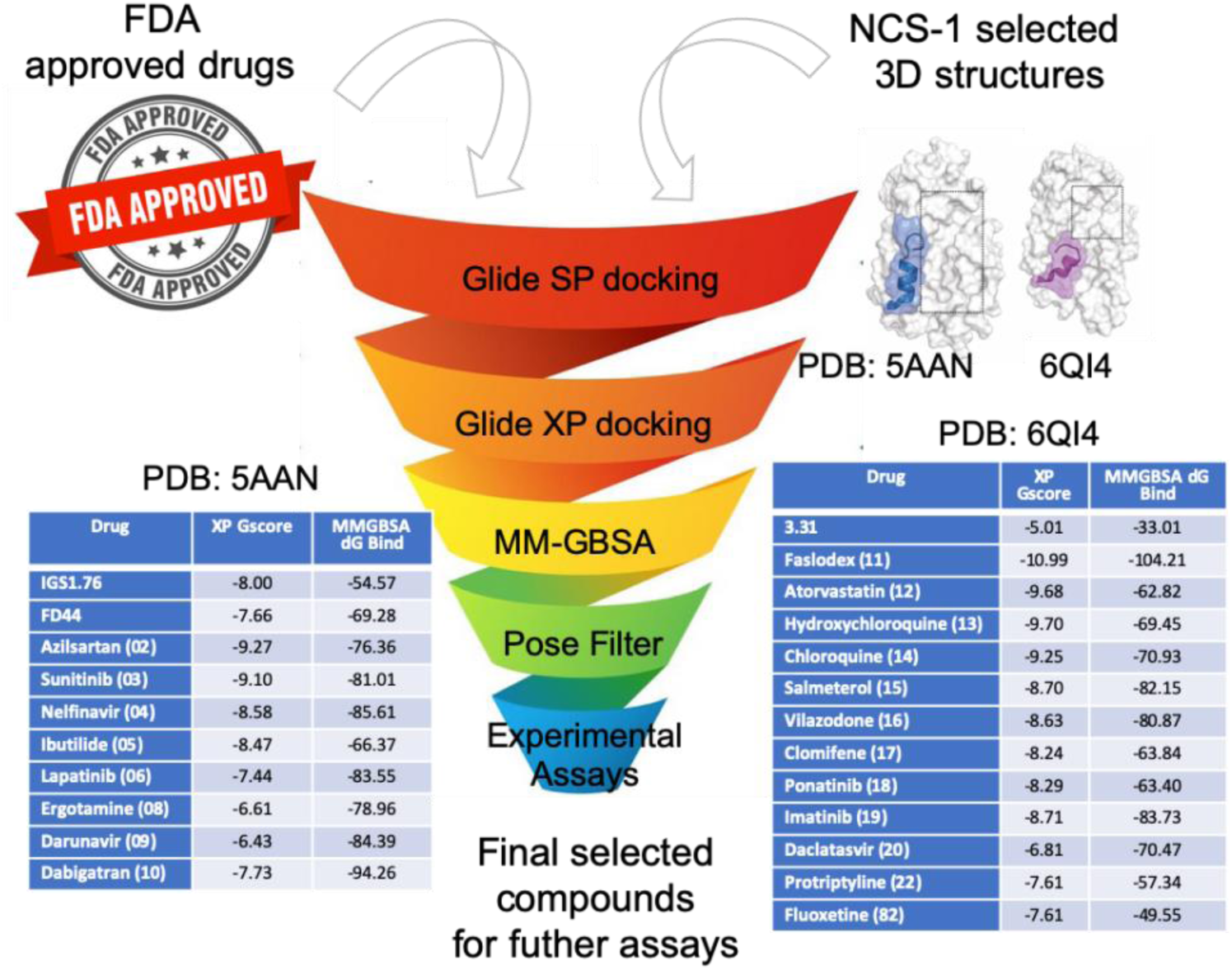
Virtual screening-selected FDA-approved compounds. Score units of XP and MM-GBSA is kcal/mol. In parenthesis, the FDA code given for each compound is shown. Control compounds are shown: IGS1.76, FD44 and 3b (21, 31, 22).

Additional filters were applied to prioritize molecules with sufficient hydrophobic character to ensure blood brain barrier permeability. Among them, compounds intended for a veterinary and/or cosmetic use, biocides, laxative or topical-administered drugs were not considered for experimental assays. Ultimately, 20 hit molecules were selected for experimental evaluation (Figure 4).

### The binding of candidate FDA compounds to NCS-1

To experimentally assess the binding of the FDA compound library to NCS-1 we used tryptophan emission fluorescence (21, 31, 22). Fluorescence intensity changes were measured at 320 nm in full-length NCS-1 upon increasing concentrations of the selected compounds (Figure 5A). Compounds exhibiting fluorescence emission within the protein’s emission range (azilsartan medoxomil (FDA02), darunavir (FDA09), faslodex (FDA11), salmeterol (FDA15), ziprasidone (FDA21) and protriptyline (FDA22)) were further analyzed using surface plasmon resonance (SPR) (Figure 5B). In these experiments, full-length NCS-1 was crosslinked to the surface of the biosensor, through its lysine amino acids, since these residues are concentrated on the upper surface and opposite to the hydrophobic crevice. This arrangement ensured that the hydrophobic crevice remained exposed to the solvent.

**Figure 5:**
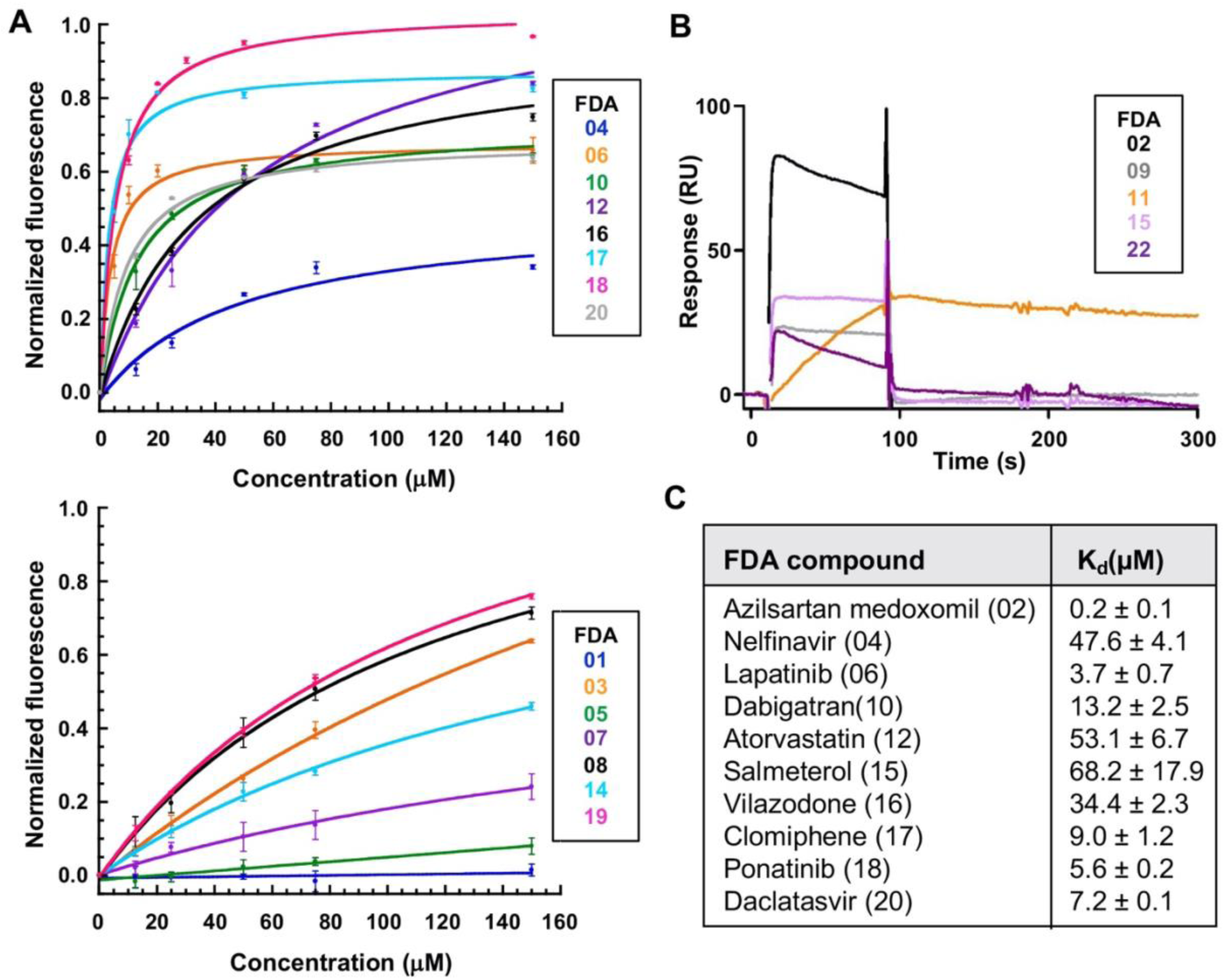
The binding of FDA hits to NCS-1: **(A)** Trp emission fluorescence assays. The upper graph displays compounds with affinities below 100 µM, while the lower graph shows those with affinities above 100 µM. **(B)** Surface plasmon resonance experiments. Binding of the tested compounds at 100 µM over CM4-immobilized NCS-1. Each compound is indicated with a different color code. **(C)** The binding affinities of the best FDA-selected compounds. Dissociation constants (K_d_ ± SD) are indicated. Only compounds with a K_d_ below 100 µM are shown. In parenthesis, the FDA code given for each compound is shown.

Due to the low solubility of ziprasidone (FDA21) under the experimental conditions, it was not possible to obtain affinity data and the compound was discarded. Compounds that failed to bind NCS-1 or exhibited a dissociation constant (K_d_) above 100 µM were excluded from further analysis. This group included amikacin (FDA01), sunitinib (FDA03), ibutilide (FDA05), capastat (FDA07), ergotamine (FDA08), hydroxychloroquine (FDA13), chloroquine (FDA14), imatinib (FDA19), fluoxetine (FDA82) and protriptyline (FDA22) (Figure 5). In contrast, compounds with affinities below 100 µM, including azilsartan medoxomil (FDA02), nelfinavir (FDA04), lapatinib (FDA06), dabigatran (FDA10), atorvastatin (FDA12), salmeterol (FDA15), vilazodone (FDA16), clomiphene (FDA17), ponatinib (FDA18) and daclatasvir (FDA20), were selected for further evaluation or their potential to modulate protein-protein interactions (Figure 5C).

### The PPI modulatory activity of the FDA compounds

To identify the FDA-approved compounds capable of disrupting the interaction in physiological conditions we have used the Proximity Ligation Assay (PLA) technology, that allows the evaluation of protein-protein interactions *in situ* with high sensitivity (32). This tool is based on the specific antibody recognition of the two proteins of interest and takes advantage of DNA primers covalently linked to the secondary antibodies. Only when the complex is formed, probe primers bind, amplification occurs, and fluorescent detection is possible.

The extent of NCS-1/D_2_R interaction was quantified after 16 hours of treatment with 5 µM of the selected FDA-approved compounds. Data are normalized to vehicle-treated cells (set to 100%). Only three compounds: azilsartan medoxomil (FDA02), atorvastatin (FDA12) and vilazodone (FDA16) had the majority of values below the 100% threshold, highlighting their inhibitory potential (Figure 6A).

**Figure 6.**
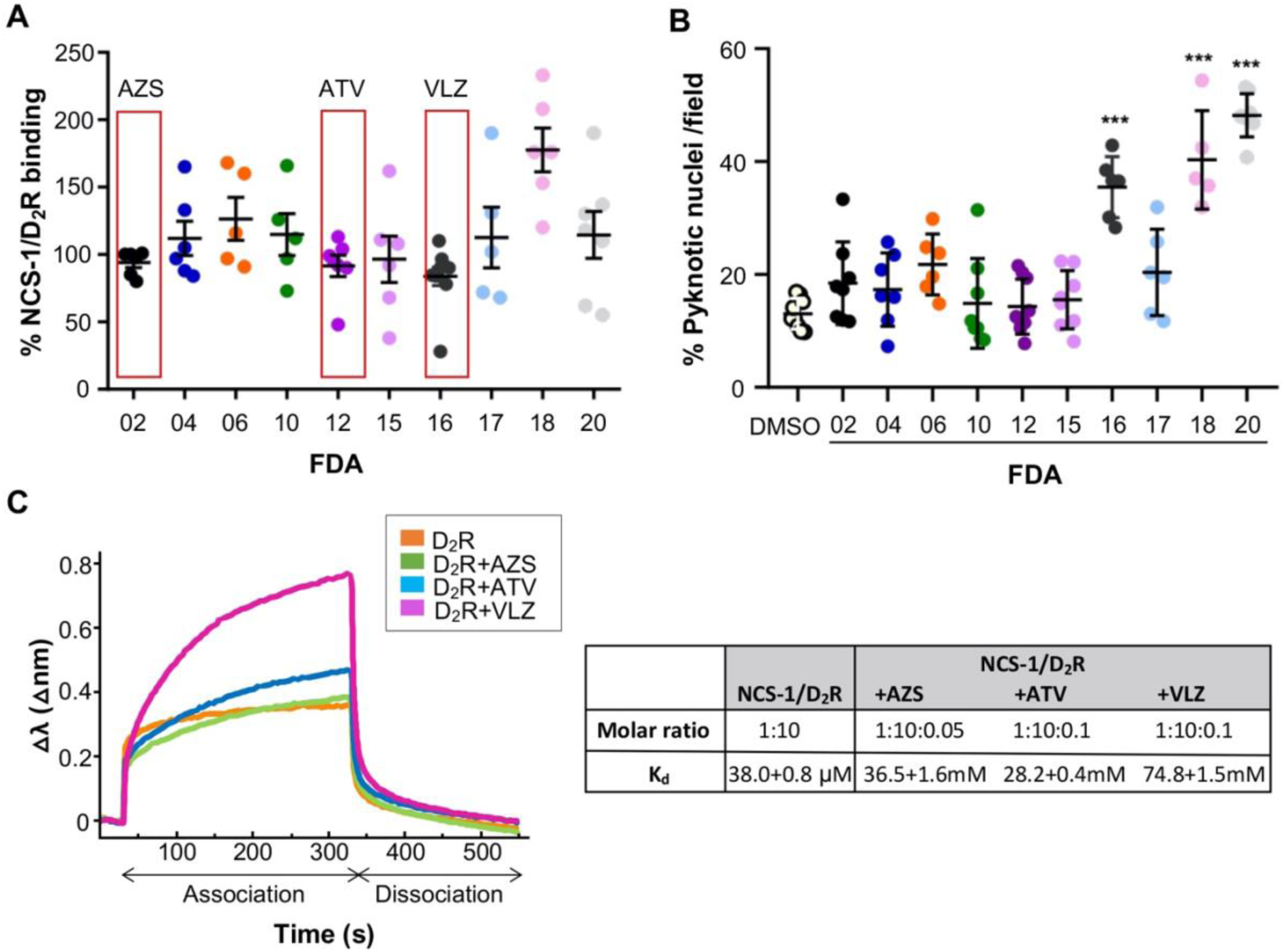
Effect of FDA-approved compounds on NCS-1/D_2_R interaction and cellular toxicity. **(A)** Proximity Ligation Assay (PLA) assessing the interaction between NCS-1 and D_2_R in HEK293 cells treated for 16 hours with 5 µM of the indicated FDA-approved compounds. Data are expressed as a percentage of NCS-1/D_2_R binding relative to vehicle-treated controls (set at 100%). Each dot represents an independent experiment; horizontal lines indicate mean ± SD. Red boxes highlight compounds for which the majority of values fall below 100%, indicating potential inhibition of the NCS-1/D_2_R PPI. **(B)** Cytotoxicity analysis of the same compounds (5 µM, 16-hour treatment) in HEK293 cells, assessed by the percentage of pyknotic nuclei per field. Data represent two fields per condition from three independent experiments. DMSO was used as a vehicle control. Bars represent mean ± SD. ***p < 0.001. **(C)** The *in vitro* binding between His-NCS-1 and a D_2_R H8 peptide (25) in the absence or presence of azilsartan medoxomil (AZS), atorvastatin (ATV) and vilazodone (VLZ). Representative biolayer interferometry sensograms showing association and dissociation of the D_2_R peptide over the time. Data are represented as the wavelength shift (Δλ) during the association and dissociation phases. A table summarizing the molar ratio (NCS-1:D_2_R H8:FDA) and the apparent K_d_ values (mean ± SEM) between NCS-1 and D_2_R H8 in the absence or presence of ligands is shown.

To exclude cytotoxic effects as a confounding factor, we performed a nuclear morphology-based toxicity. The percentage of pyknotic nuclei was quantified in HEK293 cells treated with the same compounds under identical conditions. The majority of compounds did not show significant cytotoxicity compared to vehicle (DMSO) treated cells. Although cells treated with vilazodone (FDA16) exhibited significantly higher toxicity compared to vehicle-treated cells, cell death did not exceed 50% (Figure 6B). Due to its stronger inhibitory effect on the NCS-1/D_2_R interaction, further studies were pursued with vilazodone alongside atorvastatin and azilsartan medoxomil.

To further evaluate the inhibitory activity of azilsartan medoxomil, atorvastatin and vilazodone on NCS-1/D_2_R PPI, biolayer interferometry (BLI) assays were performed using a peptide containing the D_2_R helix H8 (25). The affinity of NCS-1 for the D_2_R peptide was analyzed in the presence and absence of these compounds, as the binding of such small molecules are not directly detectable in single-channel BLItz systems. In the absence of FDA compounds, the measured NCS-1/D_2_R affinity was consistent with previous binding studies, showing an apparent affinity of 38±0.8 µM (Figure 6C) (25). However, the addition of the FDA molecules significantly weakened the PPI, shifting the dissociation constant from the micromolar to the millimolar range (Figure 6C). Since azilsartan is the molecule with higher affinity for NCS-1 (Figure 5), lower amounts of compound are needed to achieve inhibition.

### In vivo activity of the FDA-approved NCS-1/D_2_R inhibitors

As we have demonstrated thus far, the primary role of NCS-1 is to regulate the amount of D_2_R available at the plasma membrane, a function of critical importance, as it directly influences the activity of the postsynaptic neuron, or of dopaminergic neurons in cases where D_2_R acts as an autoreceptor. Therefore, if the compounds identified and characterized in this work as inhibitors of the NCS-1/D_2_R interaction are effective, they should interfere with the role of NCS-1 to promote D_2_R trafficking to the membrane. To test this hypothesis, D_2_R localization experiments were repeated in cells co-transfected with D_2_R and NCS-1 in the presence of the compounds. The results revealed a complete reversal of the effects induced by NCS-1 overexpression (Figure 7). Prior to this, it was confirmed that compound treatment in the absence of NCS-1 overexpression had no impact on D_2_R localization (Supplementary Figure 2), indicating that the observed effects are specifically mediated through disruption of the NCS-1/D_2_R interaction.

**Figure 7:**
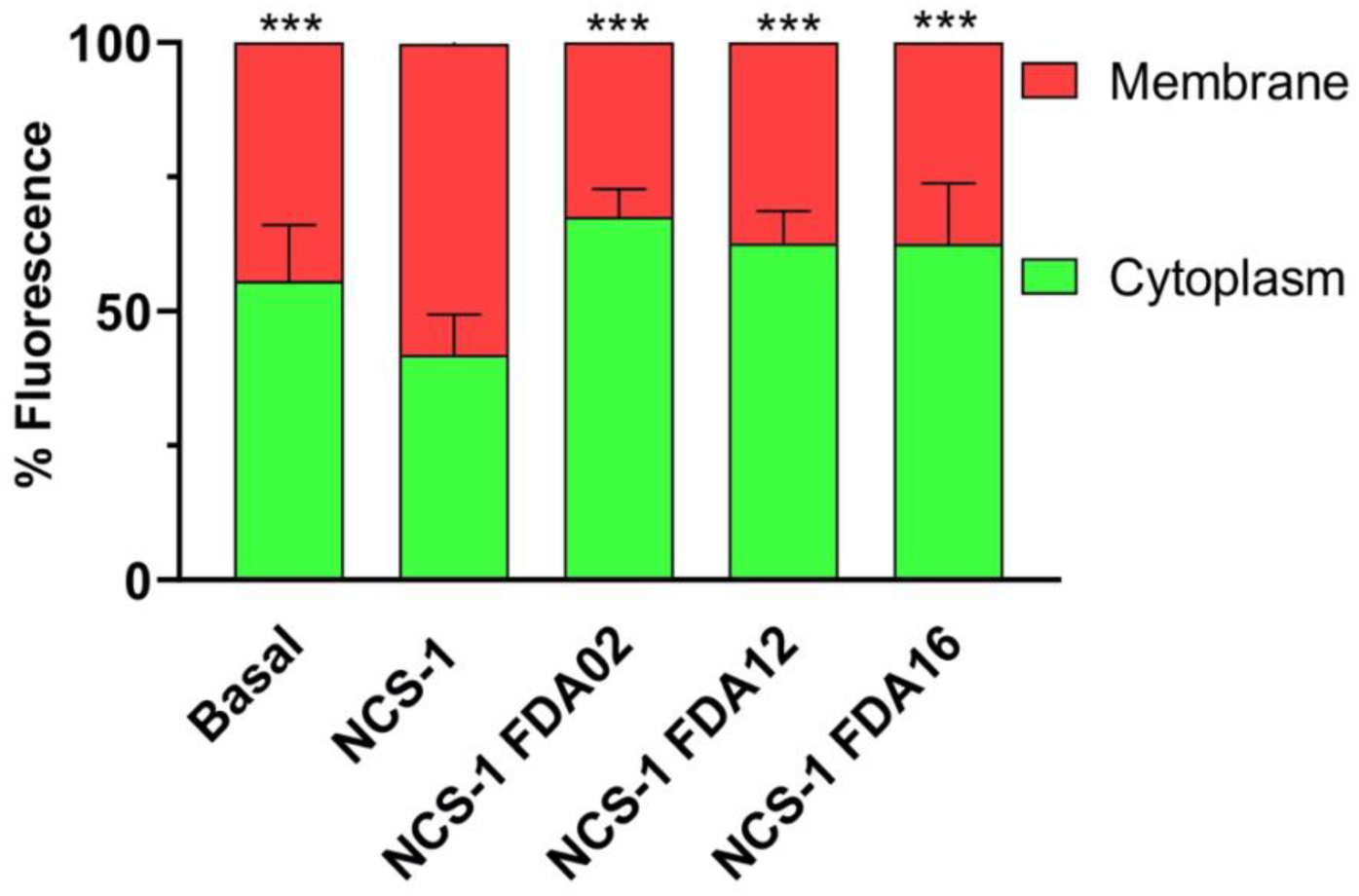
Effect of the compounds on NCS-1 mediated D_2_R cellular localization. Quantification of D_2_R fluorescence distribution between the plasma membrane (red) and cytoplasm (green) in HEK293 cells transfected with D_2_R alone or co-transfected with NCS-1, with or without a 2-hour treatment using 5 µM of the indicated FDA compounds. Statistical comparison was performed with cells cotransfected with NCS-1 without compounds. Bars represent the mean ± SD of five fields per condition from three independent experiments. ***p < 0.001.

### The mechanism of action of the FDA-approved NCS-1/D_2_R inhibitors

To investigate the mechanism by which azilsartan medoxomil (AZS), atorvastatin (ATV), and vilazodone (VLZ) modulate the NCS-1/D_2_R protein-protein interaction, the crystal structures of NCS-1 in complex with these compounds were determined. Crystals were obtained for full-length NCS-1 in complex with ATV and NCS-1ΔH10 in complex with AZS and VLZ (see Materials and Methods). The structures were solved by molecular replacement using the structure of NCS-1 (PDB code: 6QI4; (22)).

### The structure of the NCS-1/AZS complex

The asymmetric unit contained a single molecule of NCS-1ΔH10 (Table 1). However, the 2F_o_-F_c_ and the F_o_-F_c_ electron density maps revealed domain swapping between EF-hand motifs, with two NCS-1 molecules exchanging their EF-hand EF-4 (Figure 8A and Supplementary Figure 3A). The resulting NCS-1 structure, comprising EF-hands 1 to 3 from one molecule and EF-hand 4 from a second molecule, closely resembles the canonical monomeric conformation in which EF-hands pair in a two-by-two arrangement without domain swapping. The root-mean-square deviation (RMSD) for Cα atoms is 1.2 Å when compared to the reference NCS-1 structure (PDB ID: 6QI4; Canal-Martin et al., 2019).

**Figure 8:**
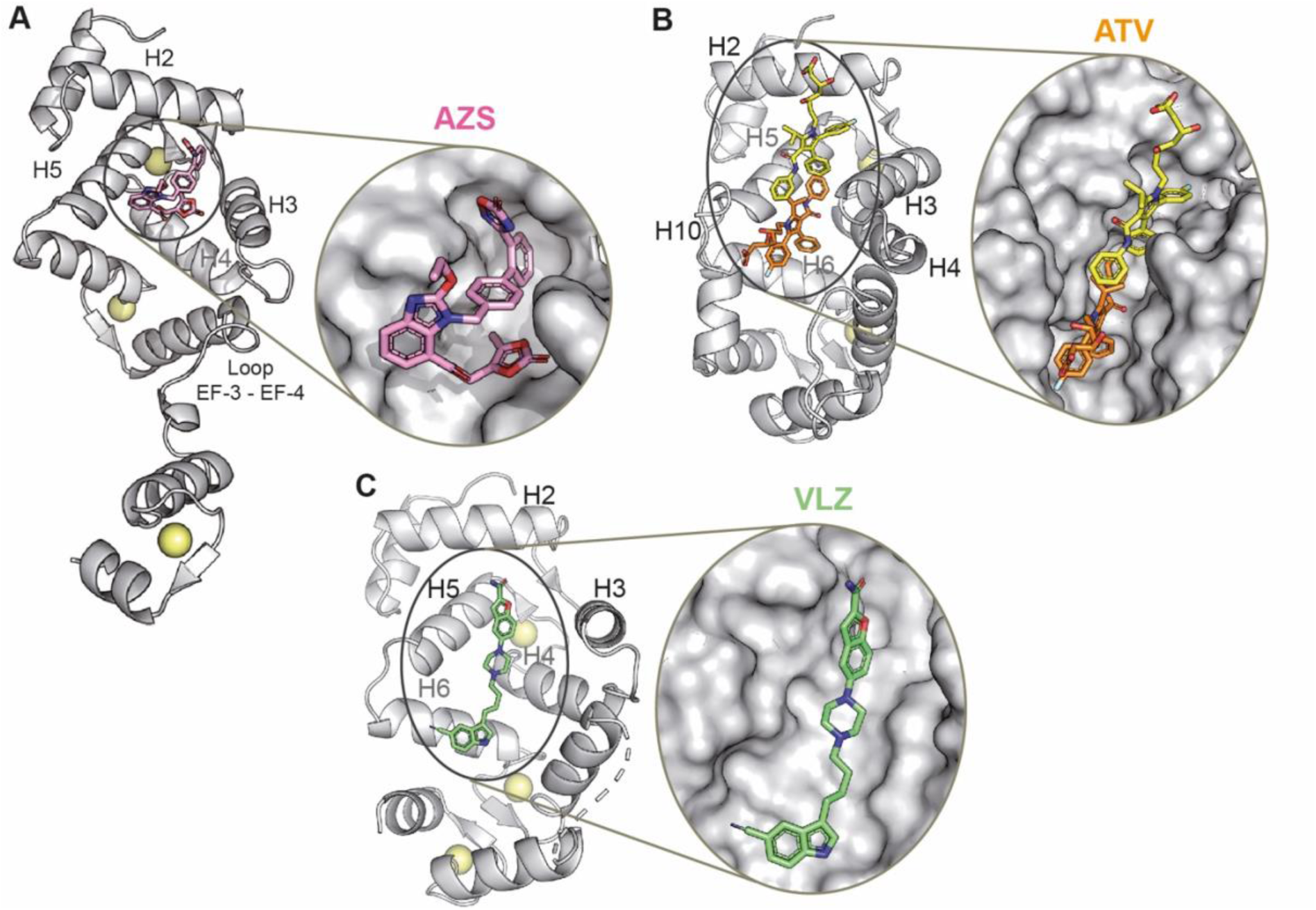
The crystal structures of NCS-1 in complex with the D_2_R PPI inhibitors. **(A)** Azilsartan medoxomil (AZS), **(B)** atorvastatin (ATV) and **(C)** vilazodone (VLZ) are shown in stick mode, NCS-1 in grey ribbons and Ca^2+^ ions as yellow spheres. A zoom on the ligand binding site is shown. NCS-1 molecular surface is represented. The NCS-1 helices participating in the recognition are indicated.

**Table 1:**
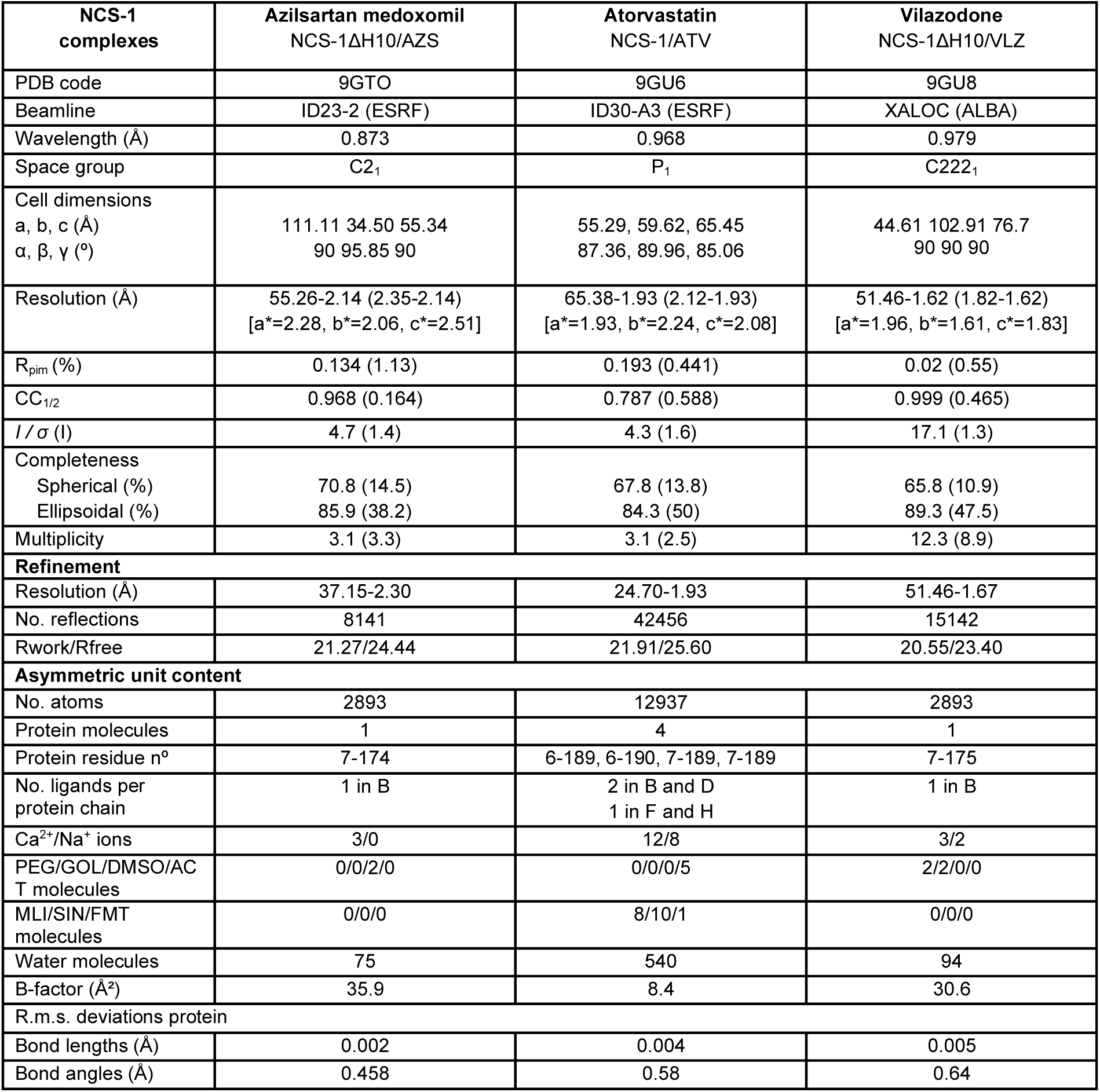
Processing and refinement statistics of NCS-1/FDA crystals.

A single molecule of AZS was bound to NCS-1, occupying the upper region of the protein’s hydrophobic cavity (Figure 8A and Supplementary Figure 3B-C). The interface between AZS and NCS-1 spans a contact area of 468.8 Å², comprising 490 interactions, predominantly mediated by van der Waals forces. Ligand recognition involves fifteen NCS-1 residues located within EF-hand 1 (helices H2 and H3) and EF-hand 2 (helices H4 and H5). The carbonyl oxygen of the AZS ester moiety forms a direct hydrogen bond with Y52, positioned at the base of the cavity. Additionally, AZS establishes two further hydrogen bonds via water-mediated interactions. Notably, all aromatic moieties of AZS—including the benzimidazole, two phenyl rings, oxadiazole, and dioxolane—engage in π-π stacking with surrounding aromatic residues (Figure 8A and Supplementary Figure 3C).

Since NCS-1 crystallized in the presence of AZS as a domain-swapped dimer, molecular dynamics (MD) simulations were performed with monomeric full-length NCS-1 in complex with azilsartan medoxomil. A crystal structure of NCS-1 in which the helix H10 is located outside the hydrophobic cavity was used (PDB ID: 6QI4; (22)). AZS coordinates from the NCS-1/AZS crystal structure (Figure 8A) (PDB code: 9GTO) were transferred to the full-length NCS-1 structure. Following structural alignment and coordinate transfer, a 1000 ns molecular dynamics (MD) simulation was performed. Throughout the simulation, a rearrangement of helix H10 was observed and after approximately 500 ns, the helix inserted into the hydrophobic crevice (Movie 1) and remained in that conformation for the rest of the simulation, as indicated by the RMSD values of the backbone atoms (Supplementary Figure 4A).

To identify the most representative and stable structure from the MD simulations, a clustering analysis was performed (Supplementary Figura 4B). A total of ten clusters were initially identified using a hierarchical agglomerative (bottom-up) algorithm with average-linkage, which considers the average distance between members of two clusters. Among the resulting clusters, Cluster 0 emerged as the most populated, comprising 42.8% of the total frames (42,764 frames). This cluster exhibited an average intra-cluster distance of 1.60 Å with a standard deviation of 0.32 Å, indicating a relatively compact conformational ensemble. The centroid of this cluster corresponds to frame 68,966, which was selected as the representative structure for further analyses. Additionally, the average distance of the cluster members to the centroid (AvgCDist) was 2.54 Å, further supporting the centrality of the selected representative conformation.

Cluster 0 appeared after approximately 50,000 frames (corresponding to 500 ns), which coincides with the stabilization of the structure. The superposition of the representative NCS-1/AZS complex of Cluster 0 with the crystal structure yields an RMSD of 1.48 Å, indicating a close agreement between the simulated and experimental binding modes (Figure 9). In this representative structure, a hydrogen bond is formed between a carbonyl oxygen from AZS and the backbone nitrogen from L189, which is located at the end of helix H10 (Figure 9 and Supplementary Figure 5). In addition, a weak hydrogen bond is transiently formed between an oxygen from the AZS oxadiazole group and the F85 side chain. Analysis of the binding site revealed that the ligand is in close proximity to 16 residues, forming van der Waals contacts. These residues are the same as those described in the crystal structure with the exception of L189 (Supplementary Figure 3C).

**Figure 9:**
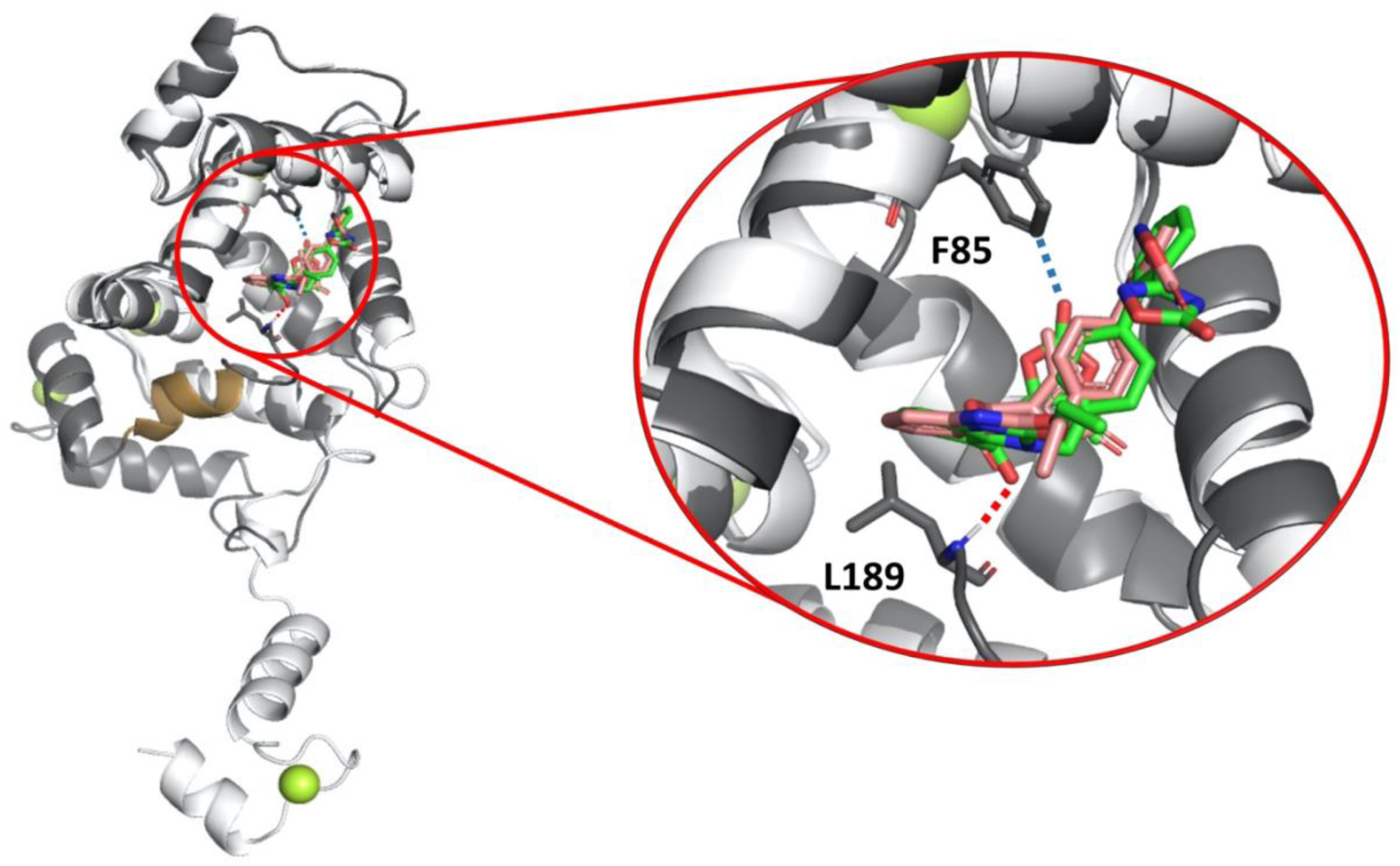
Molecular dynamics simulations of the NCS-1/AZS complex. Superposition of the crystal structure (PDB ID: 9GTO) and the representative structure from cluster 0 obtained from molecular dynamics (MD) simulations. The X-ray crystal structure is shown in light gray (AZS in pink), while the MD-derived structure is depicted in dark grey (ligand in green). Helix H10 is highlighted in gold. A close-up view of the AZS binding mode is also provided.

### The structure of the NCS-1/ATV complex

The crystal structure of NCS-1 in complex with atorvastatin revealed four independent NCS-1 molecules in the asymmetric unit (Table 1 and Supplementary Figure 6A-C). The 2Fo–Fc and Fo–Fc electron density maps showed NCS-1:ATV complexes with both 1:1 and 1:2 stoichiometries, where the drug occupied the hydrophobic crevice of the calcium sensor (Figure 8B and Supplementary Figure 6A-C). NCS-1 molecules B and D contained two atorvastatin molecules: the first (ATV1) binds near the entrance of the crevice, while the second (ATV2) is positioned deeper within the cavity (Figure 8B and Supplementary Figure 6A-B). In contrast, NCS-1 molecules F and H contained only one ligand (ATV1), located at the same site as in molecules B and D. Differences among the four NCS-1 molecules are limited to the ligand-binding regions, and within each pair (B/D and F/H), the protein structures are virtually identical, with RMSD values for Cα atoms of 0.151 Å (B vs D) and 0.155 Å (F vs H). Comparisons between molecule D and F or H yields RMSDs of 0.795 or 0.752 Å, respectively.

Within the asymmetric unit, molecules B and F interact through their Ca²⁺-binding interfaces, while molecules D and H associate via the opposite face, which includes the hydrophobic crevice (Supplementary Figure 6A). Chains D and H are arranged antiparallel to each other (Supplementary Figure 6B). Notably, in both cases, ATV1 is buried within the protein interface and shielded from solvent by an adjacent NCS-1 molecule, with F58 acting as a lid. In contrast, the second ligand, ATV2, remains solvent-exposed (Supplementary Figure 6B). All ATV1 and ATV2 molecules adopt similar conformations (Supplementary Figure 6D-E). However, weak or discontinuous electron density was observed for the 3,5-dihydroxypentanoyl tail, indicating flexibility and weak interactions with NCS-1. This region displayed the highest conformational variability. The most notable difference among the ATV2 ligands lies in the benzamide group, which interacts with adjacent ATV1 molecules. While variable in chains F and H, this group adopts a stable conformation in chains B and D to interact with ATV2 (Supplementary Figure 6C-E).

Protein-ligand interaction analysis has been made on NCS-1 chain D, where electron density for both ATV ligands was best (Supplementary Figure 6C). Both ligands span from the very top to the middle of the hydrophobic crevice. ATV1 engages 10 NCS-1 residues in helices H2, H3, and H5 (Figure 8B and Supplementary Figure 6F,H), forming 355 contacts across a 423 Å² interface. Binding is dominated by van der Waals interactions, with aromatic π–π stacking playing a major role. The central pyrrole ring of ATV1 is sandwiched between aromatic side chains on both sides of the cavity. Additional hydrophobic contacts further stabilize the complex. The benzamide moiety of ATV1 also interacts with the methylpropane group of ATV2 (Supplementary Figure 6F-I). ATV2 interacts with 13 residues (Supplementary Figure 6G,I). Its central pyrrole ring lies between T92 and Y52, both of which form direct hydrogen bonds with the adjacent amide group. A hydroxyl group on the dihydroxypentanoyl tail forms a water-mediated hydrogen bond with T92 and contacts the C-terminal end of helix H10. Additionally, the pyrrole nitrogen forms an H-bond with a malonate ion present in the crystallization solution. Although ATV2 also establishes van der Waals contacts with aromatic and hydrophobic residues, it shows fewer π–π interactions than ATV1, and the distances between its aromatic moieties is greater (Supplementary Figure 6H-I).

### The structure of the NCS-1/VLZ complex

The 2Fo–Fc and Fo–Fc electron density maps revealed a single copy of vilazodone bound within the hydrophobic groove of NCS-1. The ligand is very linear, compared with VLZ and ATV and binds the upper and central regions of the groove (Table 1, Figure 8C and Supplementary Figure 7A). The NCS-1–vilazodone interface spans 420 Å² and involves 396 contacts across 13 NCS-1 residues (Supplementary Figure 7B-C). Vilazodone primarily engages the left side of the cavity, interacting with residues from helices H2, H5, and H6 (Figure 8C and Supplementary Figure 7). Additional van der Waals interactions are contributed by the lower part of VLZ, involving helices H4 and H6. A key interaction is a hydrogen bond between Y108, located at the bottom of the binding pocket, and the indole ring of vilazodone. On the opposite side of the cavity, the ligand’s amide group forms a hydrogen bond with a polyethylene glycol (PEG) molecule. In the central region, the nitrogen atom of the piperazine ring forms an H-bond with a water molecule. Furthermore, extensive van der Waals contacts are observed with surrounding hydrophobic and aromatic residues, including notable π–π stacking between the benzofuran moiety of vilazodone and the side chain of F72 (Figure 8C and Supplementary Figure 7).

## DISCUSSION

Targeting protein–protein interactions (PPIs) remains a challenging but promising strategy in drug discovery, particularly for neurological disorders where modulating function without full inhibition can be therapeutically advantageous. Here, we focused on disrupting the interaction between the calcium sensor NCS-1 and the dopamine D2 receptor, a PPI that regulates receptor localization without altering its canonical signaling profile. This approach differs fundamentally from direct receptor antagonism, offering a way to influence dopaminergic tone by altering receptor availability at the membrane rather than interfering with ligand binding or G protein coupling. By biasing the receptor toward intracellular pools and away from the presynaptic membrane, dopaminergic neurons can attenuate the autoinhibitory feedback normally mediated by surface D_2_ autoreceptors, thereby sustaining physiologically appropriate dopamine release without changing ligand affinity or downstream signaling capacity. While reducing D_2_R presence at the presynaptic membrane may enhance dopamine release, this redistribution could also affect postsynaptic signaling, particularly in striatal medium spiny neurons (MSNs) expressing D_2_Rs. It is conceivable that any reduction in postsynaptic D_2_R activity could be counterbalanced by increased extracellular dopamine resulting from presynaptic disinhibition. Further in vivo studies will be required to determine the net effect of this shift in receptor distribution on dopaminergic circuit function.

NCS-1 has been proposed to stabilize D_2_R at the plasma membrane by blocking internalization, possibly through inhibition of GRK2-mediated phosphorylation of the D_2_R intracellular C-terminal region and interaction with GRK2 (14). However, our data suggest a different mechanism. Pharmacological inhibition experiments indicate that NCS-1 does not primarily prevent endocytosis but rather promotes D_2_R surface localization via a Brefeldin A– sensitive exocytic pathway (Figure 2). While further studies are needed to fully resolve this mechanism, our findings point toward a role for NCS-1 in facilitating forward trafficking rather than acting solely as a retention factor. Additionally, over-expression of NCS-1 does not have an impact on the pharmacological profile of the D_2_R in a cellular model system (Figure 3), concluding that NCS-1 likely interacts with D_2_R during trafficking. The change in receptor expression is however likely to change the signaling in vivo where receptor reserve has an impact on the functional outcome.

The identification of active molecules through a structure-based virtual screening of FDA-approved drugs (Figure 4) highlights the potential of drug repurposing in neuropharmacology. Among the hits, azilsartan medoxomil, atorvastatin, and vilazodone showed measurable affinity for NCS-1 (Figure 5) and disrupted the NCS-1/D_2_R interaction in vitro and in cells (Figure 6). These compounds, with documented clinical use and varying CNS penetrance, offer attractive starting points for further development.

Crystallographic and biophysical data revealed that these molecules bind within the hydrophobic crevice of NCS-1, overlapping with the D_2_R recognition site and thus competing with D_2_R for NCS-1 binding (Figure 6 and 8). Notably, the ligands exploit a combination of van der Waals contacts, π–π stacking, and hydrogen bonding, providing a diverse interaction footprint across the crevice. Azilsartan medoxomil binds to the upper region of the NCS-1 crevice, and molecular dynamics simulations suggest that its binding does not interfere with the insertion of the dynamic helix H10 into the lower part of the crevice (Figure 9). In contrast, atorvastatin and vilazodone also localize to the upper half of the crevice but extend further into the central region. These small molecules partially overlap with an inserted helix H10 (Supplementary Figure 9), suggesting that the compounds disrupt the dynamics of this critical structural element, thereby hindering its proper insertion into the binding cavity. In fact, the deletion of the helix H10 improves their binding to NCS-1 (Supplementary Figure 8). This disruption of the dynamics of helix H10 may account for the observation that, compared with AZS, both ATV and VLZ exhibit comparable inhibition of the protein-protein interaction despite their lower binding affinities (Figure 5), as ligand-induced conformational rearrangements likely destabilize the D_2_R recognition interface. These structural insights not only validate the mechanism of inhibition but also offer a template for rational optimization of NCS-1-targeted PPI modulators.

Collectively, our results reveal a novel therapeutic strategy based on modulating intracellular protein–protein interactions to regulate dopamine D_2_ receptor trafficking. Rather than blocking receptor signaling directly, this approach shifts receptor localization to fine-tune dopaminergic tone, offering a targeted alternative to conventional antagonism. Critically, the identification of FDA-approved compounds as inhibitors of the NCS-1/D_2_R interaction underscores the untapped potential of drug repurposing in neuropharmacology. While further studies are needed to determine the therapeutic relevance, pharmacological manipulation of receptor trafficking may offer a promising strategy to address dopaminergic dysregulation in conditions such as schizophrenia, bipolar disorder, and Parkinson’s disease.

## MATERIALS AND METHODS

### Virtual screening of drug-like compound

#### Ligand Preparation

An initial list of 1352 FDA-approved drugs was retrieved from the ZINC database. The conversion from SMILES to SD format was performed using the structconvert tool in the Schrödinger module (33). Ligand preparation was carried out using the LigPrep and Epik modules of the Maestro suite (34–37). Progressive levels were generated, encompassing possible ionization states at physiological pH and potential tautomers. Final energy minimization was performed using the OPLS4 force field, with default parameters set for stereoisomer (38).

#### Protein Preparation

NCS1 proteins, human NCS-1 (hNCS-1, PDB code 6QI4, (22)) and Drosophila homologue of human NCS-1 (dNCS-1, PDB code 5AAN, (21)) were prepared for subsequent computational analyses using the Protein Preparation Workflow (39–42), a tool integrated into Maestro (37). This process began with protein structure preprocessing, including bond order assignment and structural adjustments facilitated by Prime (41, 43, 44). Furthermore, PROPKA (45) was employed to refine the protein’s hydrogen bond network and determine residue protonation states at pH 7.2, leveraging its pKa prediction capabilities. Selection of the most favorable protonation state for each was based on hydrogen bond formation and an associated penalty score. The entire preparation concluded with a final restrained minimization utilizing the OPLS4 force field (38).

#### Structure-Based Virtual Screening (SBVS) and MM-GBSA Rescoring

A set of 2143 molecular candidates from the post-processed FDA database was screened against the previously described targets. The centroid of the crystallized ligand in the catalytic pocket served as the grid center. SBVS was then performed using a pipeline that included 3 stages. The first one consisted of massive docking simulations employing the Glide software (46–49) and the Standard Precision (SP) method. In this first stage, an enhanced sampling approach was used, and 5 poses were generated per compound state. The top 50% of compounds (according to the scoring function) were retained and used for the second stage, where the Extra Precision (XP) method was employed. In the second stage, 25% of the top-ranked solutions were retained. Rescoring was performed in the third stage with the Prime MM-GBSA method (41, 43, 44).

#### Docking Validation

Docking studies were validated by root-mean-square deviation (RMSD) values obtained from the superposition of the redocked ligand structure (IGS1.76) and X-ray crystal structure. The docking grid was centered on the geometric center of the ligand observed in the crystal structure within the catalytic pocket.

### Molecular dynamics simulations

The movement of helix H10 towards the crevice and the stability of the X-ray structure NCS-1/AZS was studied using MD simulations with AMBER20 (50). The NCS-1 structure was prepared using Protein Preparation Workflow (39–42), was modified to create zero-order bonds for calcium ions, and all water molecules were removed. The protein model was constructed using the ff14SB force field (51) with the TIP3P water model (52). Three calcium ions were maintained, and neutralizing sodium counterions were added as needed to ensure system neutrality.

Energy minimization was performed to relieve steric clashes and optimize geometry. The protocol consisted of five sequential steps. The first four steps involved positional restraints on the protein backbone with gradually decreasing force constants. These restrained minimization phases employed a hybrid steepest descent/conjugate gradient algorithm, a non-bonded cutoff of 10.0 Å, and a final RMSD convergence criterion of 0.005 Å. Following energy minimization, molecular dynamics (MD) simulations were performed to equilibrate the protein-ligand complex in explicit solvent. The system was thermalized and equilibrated using a series of simulations: initial heating to 100 K with protein restraints (10 ps), followed by further heating to 300 K with restraints (100 ps), and a final 100-ps production run at 300 K without restraints. Temperature and pressure control were achieved using a Langevin thermostat (damping coefficient of 1.0 ps-1) (53, 54) and a Berendsen barostat (55), respectively. Bonds involving hydrogen atoms were constrained using the SHAKE algorithm (56), and a 10.0 Å cutoff was applied for non-bonded interactions. Production simulations were performed at 300 K under constant pressure and temperature. To generate a 1000-ns trajectory, a 1-ns production protocol was repeated 1000 times. The cpptraj module of AMBER was used for trajectory analysis (57).

### Molecular cloning and constructs

Human NCS-1 constructs, both untagged and V5-tagged, were previously described (58, 21). The cDNAs of the human D_2_R (HASS-FLAG-EGFP-3C_protease_-D_2_R), MeN (NanoLuc Fragment-membrane anchor) and β-Arrestin-2 (β-Arrestin-2-NanoLuc Fragment) were obtained through gene synthesis (Gene Fragments, Twist Bioscience) and cloned into the pCDNA4 using in vivo DNA assembly (59, 60). D_2_R was subsequently subcloned into the pCDNA3.1/V5-His-TOPO vector for proximity ligation experiments. The Gα_OA_-RLuc8, Gβ_3_, and Gγ_9_-GFP2 constructs in pCDNA5 and pCDNA3.1 were a gift from Bryan Roth’s lab (Addgene plasmid kit #1000000163). GRK2 (pcDNA4) was a kind gift from Nevin Lambert. The plasmids encoding the human D_2_R linked to the natural peptide and the miniGα_O_ linked to the large-BiT were previously described (61, 62) and kindly provided by Dr. K. Sahlholm (Karolinska Institute, Stockholm, Sweden) and Dr. S. Ferré (National Institutes of Health, Baltimore, MD, USA), respectively.

For biophysical and crystallographic assays, full-length human NCS-1, His-tagged full-length NCS-1 (His-NCS-1), and an helix H10 deletion construct (NCS-1ΔH10) were expressed and purified in Ca^2+^ saturating conditions as published elsewhere (16, 22, 63).

### Cell culture and transfection

Human embryonic kidney HEK293T cells (ATCC, not authenticated in-house) were grown in DMEM supplemented with 100 U/ml streptomycin, 100 mg/ml penicillin and 10% (v/v) foetal bovine serum. Manipulation and maintenance were done in a biological safety cabinet class 1 and in an incubator at 37°C, 5% CO_2_ and 90% relative humidity. HEK293T cells were transiently transfected with the indicated cDNA construct using polyethylenimine (PEI MAX™, Polysciences Inc.) at a 6:1 PEI:DNA ratio. For D_2_R localization experiments and Proximity ligation assays, transfections were carried out using Lipofectamine 2000 (Invitrogen) in Opti-MEM (Thermo Fisher Scientific) without serum or antibiotics, following the manufacturer’s instructions. After a 4-hour incubation with the DNA-Lipofectamine mixture, the medium was replaced with a fresh complete medium, and cells were incubated for an additional 24 hours before further processing.

### Immunostaining for D_2_R localization

HEK293 cells were seeded on poly-D-lysine-coated glass coverslips in 24-well plates. Following transfection and drug treatments, cells were fixed with 2% paraformaldehyde (PFA) for 10 minutes and with 4% PFA for an additional 10 minutes at room temperature. Coverslips with fixed cells were then blocked in PBS containing 10% normal goat serum (NGS) for 30 minutes at room temperature.

To differentiate membrane-localized from cytoplasmic D_2_R, a two-round immunostaining protocol was performed. In the first round, coverslips were incubated with rabbit anti-FLAG primary antibody (Sigma, F7425; 1:1000) for 2 hours at room temperature, followed by Alexa Fluor 594-conjugated anti-rabbit secondary antibody (Thermo Fisher Scientific, 1:500) for 2 hours. After three PBS washes, fluorescence was fixed with 10% formaldehyde for 3 minutes. The second round started with a blocking step using PBS containing 10% NGS and 0.03% Triton X-100 for 1 hour at room temperature. Then and incubation with mouse anti-FLAG primary antibody (Sigma, 1:1000) overnight at 4°C, followed by Alexa Fluor 488-conjugated anti-mouse secondary antibody (Invitrogen, A10684; 1:500) for 2 hours at room temperature. Coverslips were washed with PBS and mounted using ProLong™ Gold Antifade (Thermo Fisher Scientific) with Hoechst for nuclear counterstaining.

### Proximity Ligation assay

HEK293 cells were co-transfected with plasmids encoding human NCS-1 and D_2_R tagged with V5. 24 hours post-transfection, cells were treated for 16 hours with either FDA-approved compounds or an equivalent volume of DMSO, serving as the vehicle control.

To assess protein-protein interactions, a Proximity Ligation Assay (PLA) was performed in conjunction with flow cytometry. PLA enables in situ detection of proteins in close proximity (within 40 nanometers) by employing antibodies conjugated to DNA oligonucleotides (32). Upon binding their respective targets, these probes initiate rolling-circle amplification, producing a fluorescent signal. After treatment, cells were detached using trypsin and gently dissociated then fixed for 20 minutes using the Fixation Reagent and then permeabilized with the Permeabilization Reagent, both from the Intracellular Fixation and Permeabilization Buffer Set (eBioscience, Thermo), specifically optimized for flow cytometry applications. Primary antibodies; mouse anti-V5 (1:2,000, Thermo Fisher Scientific) and rabbit anti-NCS-1 (1:1,000, Cell Signaling) were diluted in the permeabilization buffer and added for overnight incubation. Following washes, cells were incubated with Duolink PLA probes: anti-rabbit MINUS and anti-mouse PLUS (Sigma). Negative controls omitting the primary antibodies were included to assess background signal. Detection was carried out using the Duolink PLA Flow Cytometry Green Detection Kit (Sigma), according to the manufacturer’s instructions. Fluorescent signals were analyzed using a BD FACS Canto II flow cytometer with BD FACSDiva software.

### β-Arrestin recruitment assays

β-arrestin recruitment was assessed in HEK293T cells (ATCC, not authenticated in-house) using the β-arrestin membrane recruitment assay developed by Pedersen et al. (29). 50,000 cells/well were seeded in previously poly-lysined 96-well white plates. The following day, cells were transfected with D_2_R:MeN:β-Arrestin-2 (ArC):GRK2:pcDNA3.1 or D_2_R:MeN:β-Arrestin-2 (ArC):GRK2:NCS-1 at a 2:1:1:3:4 ratio. After 48 hours, the medium was replaced with 90 µL/well of 1× Hank’s balanced salt solution with 20 mM HEPES pH 7.4 and 7.5 µM Coelenterazine 400a. 10 µL of varying quinpirole concentrations were added and luminescence was measured using a CLARIOstar (BMG Labtech) after a 20-minute incubation. Data analysis was performed using GraphPad Prism 8.0.1. Data were normalized and a four-parameter logistic curve was fit into the data. Efficacy was normalized relative to D_2_R:pcDNA3.1. Data are presented as mean ± SEM of at least three independent experiments performed in technical triplicate. Source data is provided as a Source Data File.

### Nanobit miniGα protein recruitment assay

The NanoBiT assay was performed as previously described (28). HEK293T cells were transiently transfected with D_2_R-NP:miniG_O_-LgBiT:pcDNA4.1 or D_2_R-NP:miniG_O_-LgBiT:NCS-1 at a 10:1:20 ratio. After 36h, cells were harvested in 1× Hank’s balanced salt solution (Cytiva, Utah, USA) and transferred (90 μL) into a white 96 well plate (Corning™ 96-Well, Cell Culture-Treated, Flat-Bottom microplate; 80,000 cells/cm²). Subsequently, 10 µl of a 10 µM coelenterazine 400a solution was added to each well. After ten-minute incubation, end-point luminescence was determined using a CLARIOstar plate-reader (basal signal). Immediately after basal measurement increasing concentrations of quinpirole were added and luminescent signal was measured after 10 minutes. Data was expressed as % of receptor response which was calculated as indicated:

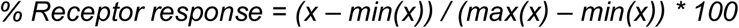

where “x” corresponds to the ratio between quinpirole and basal signal of each well and min(x) and max(x) to the ratio of the lowest and highest signal, respectively. Data analysis was performed using GraphPad Prism 8.0.1, with data normalized and fit to a four-parameter logistic curve. Results are presented as mean ± SEM from four independent experiments, each performed in triplicate.

### Cellular BRET assays

G protein dissociation was assessed using the TRUPATH system in HEK293T cells (27). 50,000 cells/well were seeded in poly-lysine coated 96-well white plates. The following day, cells were transfected with D_2_R:Gα_OA_-RLuc8:Gβ_3_:Gγ_9_-GFP2:pcDNA3.1 or D_2_R:Gα_OA_-RLuc8:Gβ_3_:Gγ_9_-GFP2:NCS-1 at a 2:1:1:1:4 ratio. After 48 hours, the medium was replaced with 90 µL/well of 1× Hank’s balanced salt solution with 20 mM HEPES pH 7.4 and 7.5 µM Coelenterazine 400a. 10 µL of varying quinpirole concentrations were added, and BRET signals were measured using a CLARIOstar (BMG Labtech) with 400 nm (RLuc8) and 498.5 nm (GFP2) emission filters at integration times of 1.85 s. BRET ratios were calculated as the ratio of the GFP2 signal to the Rluc8 signal. Data analysis was performed using GraphPad Prism 8.0.1, with data normalized and fit to a four-parameter logistic curve. Results are presented as mean ± SEM from at least three independent experiments (technical triplicates). Efficacy was normalized relative to D_2_R:pcDNA3.1. Source data is provided as a Source Data File.

### Ligand preparation

Compounds (>95 % purity) were purchased to MedChemExpress except in the case of ergotamine, darunavir and dabigatran, which were bought to Sigma. Compounds were solubilized in pure DMSO at 5 mM (biophysical studies) or 30-50 mM (crystallographic studies) with the exception of amikacin, capastat and hydroxychloroquine, which were soluble in aqueous solution.

### Protein intrinsic emission fluorescence

To determine the affinity of the FDA compounds for full-length NCS-1 or NCS-1ΔH10, we used fluorescence techniques, since the presence of tryptophan (Trp) and tyrosine (Tyr) residues in NCS-1 sequence confers the protein intrinsic emission fluorescence when excited at 295 nm. Fluorescence intensity was monitored at 330 nm and 35°C with a nano-DSF machine (nanoTemper). Protein concentration was set to 5 µM in a buffer containing Tris 50 mM pH 8.0, NaCl 125 mM, CaCl_2_ 5 µM, 5 % DMSO. Ligand concentration was increased up to 150 uM. Three independent experiments were performed for each protein:ligand ration. Previously, we verified that the compounds did not emit at 330 nm. In fact, compound FDA-02, 09, 11, 15, 21 and 22 were measured using SPR due to their high intrinsic emission at 330 nm. The apparent dissociation constant, K_d_, was obtained by using a least squares algorithm to fit the recorded data to the equation previously described. Apparent K_d_ values are reported as mean ± SD (21, 31, 16).

### Surface plasmon resonance

SPR experiments were performed at 25°C using a Biacore X-100 apparatus (Biacore, GE) with a running buffer containing Tris 50 mM, pH 8.0, NaCl 125 mM, CaCl_2_ 0.5 mM with 2% DMSO. NCS-1 was immobilized on a CM4 sensor chip (Biacore, GE) following a standard amine coupling method. The carboxymethyl dextran surface of the flow cell 2 was activated by injecting a 1:1 mixture of 0.4 M EDC and 0.1 M NHS for 7 min. The protein was then covalently coupled to the surface via a 7-min injection at several dilutions, with a final concentration of 100 µg/ml in 10 mM sodium acetate, pH 4.0. Unreacted *N*-hydroxysuccinimide esters were quenched by a 7-min injection of 0.1 M ethanolamine-HCl pH 8.0, achieving an immobilization level of 5000 RUs. Flow cell 1, subjected to the same amine coupling procedure but without protein, served as a reference. Prior to use, 10 mM stock solutions of FDA compounds were serially diluted to final concentration of 100 µM in running buffer. Typically, a series of different compounds was injected onto the sensor chip at a flow rate of 30 µl/min for 80s, followed by a 80-s dissociation rate. After dissociation, an additional wash step was performed using a 50 % DMSO solution. Regeneration was not required.

For kinetics measurements of FDA2, FDA9, and FDA15, compounds were tested at concentrations ranging from 10-100 µM. Compounds were injected at 50 µl/min for a period of 100s followed by a dissociation of 300s. Sensograms data were double-referenced and solvent-corrected using the BiaEvaluation X-100 software (Biacore, GE). FDA-21 precipitated on the sensor chip under the experimental conditions and no kinetics measurements were performed.

### Biolayer interferometry assay

A single-channel BLItz system (ForteBio) was used to assess the interaction between NCS-1 and a D_2_R H8 peptide (25), as well as its modulation by various FDA-approved compounds.

This technique immobilizes one of the molecules at the biosensor tip and analyzes interference patterns generated by white light reflection, providing an optical, label-free approach to studying macromolecular interactions. Due to the high concentration of immobilized molecules at the tip, BLI enables the detection of both low- and high-affinity binders, making it particularly suitable for studying weak interactions. Real-time shifts in the interference pattern (Δλ) occur as the number of molecules interacting with those immobilized on the biosensor tip changes. The experimental data obtained from BLI include association and dissociation rate constants, which are used to calculate the apparent equilibrium dissociation constant (Kd) (64).

Ni-NTA biosensors (Sartorius) were used to immobilize the N-terminally His-tagged NCS-1, which was prepared at 5 µM in a buffer A containing 50 mM Tris pH 8, 125 mM NaCl, 128 µM CaCl_2_, and 5 % DMSO. The immobilization of His-NCS-1 was performed in three steps: (1) baseline stabilization (buffer A, 30 s), (2) loading (His-NCS-1 in buffer A, 300 s), and (3) equilibration (buffer A, 220 s). D_2_R H8 peptide was prepared at 50 µM in buffer A. In competition assays, the 50 µM D_2_R H8 peptide solution additionally contained 0.25 µM FDA-02 or 0.5 µM FDA-12 and FDA-16. Following protein immobilization, all experiments followed the same sequence: (1’) baseline stabilization (buffer A, 30 s), (2’) association (only D_2_R H8 peptide or its mixture with FDA compounds in buffer A, 300 s), and (3’) dissociation (buffer, 220 s). Each FDA-approved compound was tested in three independent experiments. The apparent dissociation constant (K_d_) was calculated by fitting data extracted from sensograms using BLItz Pro software, assuming a 1:1 equilibrium binding model. Apparent K_d_ values are reported as mean ± SEM.

### Protein-ligand assembly, crystallization, diffraction data collection and structure solution of NCS-1 bound to FDA ligands

Prior crystallization, the contribution of the helix H10 to ligand binding was assessed in case this information could guide construct selection for structural studies. ATV displayed an improvement of the apparent binding affinity for H10-truncated NCS-1 compared with the full-length protein (53 µM vs 17 µM, respectively). Likewise, truncation of the helix H10 also enhanced VLZ binding affinity (34 µM vs 8 µM) (Figure 5C and Supplementary Figure 8). Given the initial high affinity of AZS for full-length NCS-1, the affinity for the H10-truncated construct was not measured. Initial crystallization screenings yielded to crystals of ATV in complex with full-length NCS-1. In the case of AZS and VLZ, the successful construct that yielded to crystals was the truncated NCS-1ΔH10.

Full-length NCS-1 protein was dialyzed in buffer 20 mM NaAc pH 5.5, 0.5 mM CaCl_2_, 0.5 mM DTT and 5% DMSO (2 changes, 4 and 16 h). Similarly, NCS-1ΔH10 protein was dialyzed in buffer 50 mM Tris pH 8.0, 1.75 mM CaCl_2_, 1 mM DTT and 10% DMSO. The proteins were concentrated to 15 mg/ml using a 10 kDa cut-off concentrator (Vivaspin) before ligand addition. AZS and VLZ were added to NCS-1ΔH10 in a 1:2 (protein:ligand) molar ratio, reaching a final DMSO concentration of 15% and 10%, respectively. ATV was mixed with purified full-length NCS-1 in a 1:4 molar ratio, reaching a final DMSO concentration of 5%. Finally, the protein/ligand complexes were set up for crystallization experiments.

Crystals were obtained by the co-crystallization method. After 30min protein:ligand incubation, crystallization screenings were set with an Oryx8 robot (Douglas Instruments) at 4 °C, using the sitting drop vapor diffusion method and mixing equal volumes of protein complex and precipitant. NCS-1ΔH10/VLZ complex crystallization were set up at 18 °C. Initial crystals were obtained in solutions from JBScreen Classic and JCSG++ (Jena Bioscience) and INDEX (Hampton Research) crystallization screenings. NCS-1ΔH10/AZS crystals were obtained in 70% MPD, 0.1 M HEPES pH 7.5. Prior to crystal freezing, a 3h soak with 30 mM AZS was performed. Crystals of the NCS-1/ATV complex grew using streak-seeding technique and a precipitant solution containing 70 % Tacsimate pH 7.0 at 4°C. NCS-1ΔH10/VLZ crystals grew in 20 % PEG 3000, 0.1 M HEPES pH 7.5, 0.2 M NaAc precipitant solution. Only NCS-1/VLZ crystals were cryo-protected by adding 20 % (v/v) glycerol to the crystallization solution. All crystals were flash-frozen in N_2_(l).

An NCS-1ΔH10/VLZ diffraction dataset was collected at 100 K and 0.979 Å wavelength at ALBA BL13 beamline synchrotron radiation source. NCS-1ΔH10/AZS and NCS-1/ATV diffraction data were collected at 0.873 Å wavelength at ESRF ID23-2 and ID30-A3 beamlines, respectively (Table 1). Data were processed automatically at the beamlines with AutoPROC (65). Structures were solved by the molecular replacement method with Phaser (66). The structures of hNCS-1 (PDB: 6QI4; (22)) with or without the C-terminal helix H10 were used as search models, depending on the NCS-1 construct used for crystallization. FDA ligands dictionaries with geometric restrains were generated with acedrg (67). Successive cycles of automatic refinement with Phenix (Adams et al 2010) and manual building with Coot (68) were performed. The final models were validated with Molprobity (69). Details on data processing and refinement are shown in Table 1. The structures were analyzed using different programs from the CCP4 package (70), LigPlot (71) and PISA server (72). Images were prepared with PyMOL (73). The final structures were deposited in the PDB with codes: NCS-1/AZS (9GTO), NCS-1/ATV (9GU6), NCS-1/VLZ (9GU8).

## ACKNOWLEDGMENTS

This work was funded by grants PID2019-111737RB-I00 and PID2022-137331OB-C31 (to MJSB) PID2020-113359GA-I00, PID2023-148739NB-I00 (to JGN), PID2022-137331OB-C32 (to AM), PID2022-137331OB-C33 (to NEC) and the Spanish “Ramón y Cajal” program (to JGN and AM), which were funded by the MICIU/AEI/10.13039/501100011033, FEDER and EU. SPS was supported by a contract from “Programa de Empleo Juvenil de la Comunidad de Madrid” PEJ-2020-AI/BMD-18666 and CMR. with a Spanish National Research Council JAE-Intro grant (JAEINT_23_00101). MJSB would like to thank the Biophysical Techniques Facility at IQF Blas Cabrera for access to the confocal microscope, nano-DSF and biolayer interferometry equipments; ALBA (XALOC beamline) and ESRF synchrotrons for access and support of the staff and Dr. Laura Lagartera for the SPR experiments and data analysis (Servicio de Interacciones Biofísicas, IQM-CSIC).

## Notes

### Competing Interest Statement

The authors have declared no competing interest.

