## Supplemental Figures 1 to 9 and Supplemental Table 1 for "FDA drug repurposing uncovers modulators of dopamine D_2_ receptor localization via disruption of the NCS-1 interaction"

**Supplementary Figures**

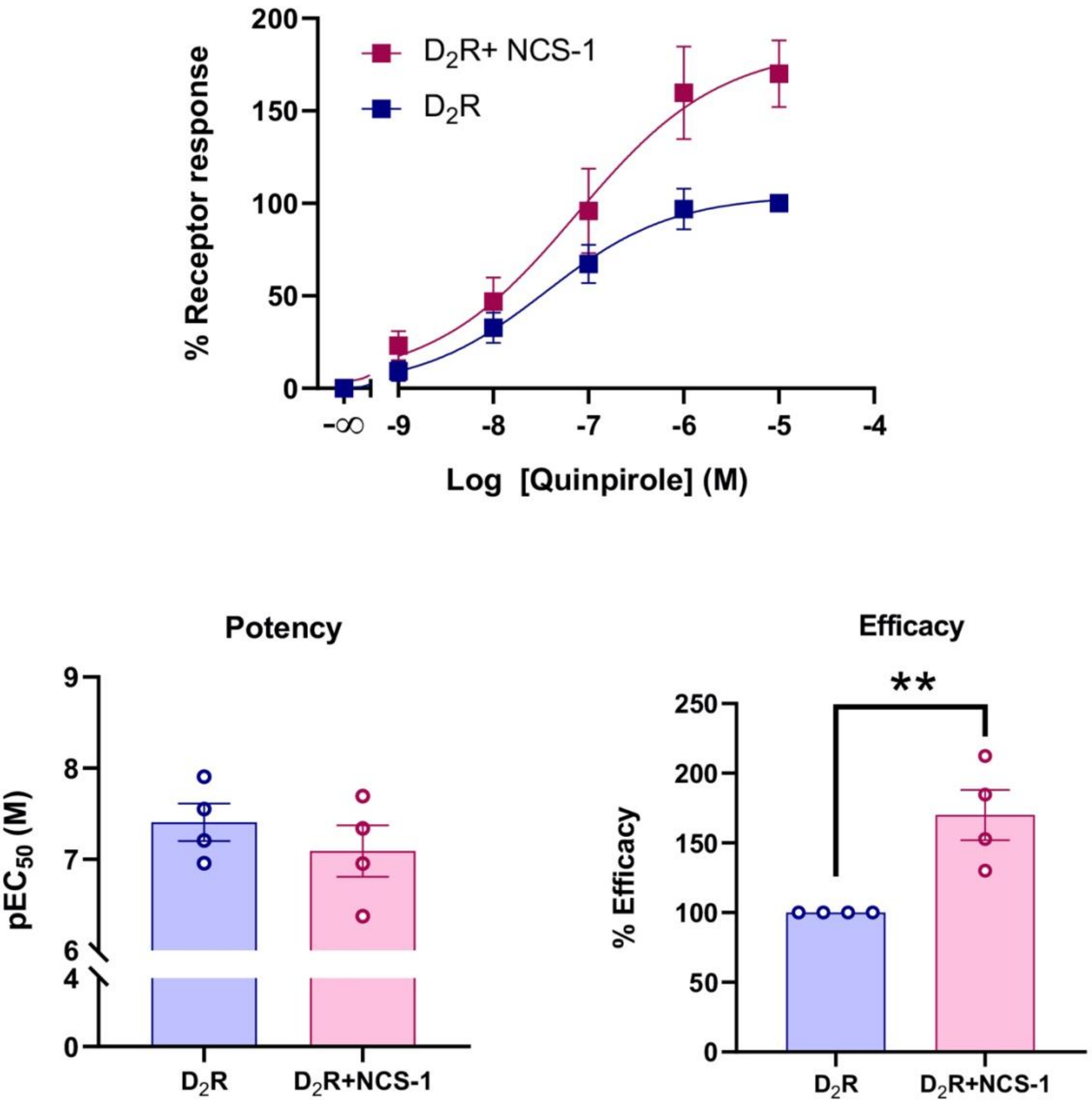


**Supplementary Figure 1: Nanobit recruitment assay for the dopamine $D_2$ receptor in the presence of NCS-1. (A)** HEK293-T cells expressing $D_2R$ linked to the natural peptide and miniGαo-LgBiT in the absence (blue squares) or presence (red squares) of NCS-1 were challenged with increasing concentrations of quinpirole and the receptor/miniG$\alpha_O$ protein coupling was determined by the NanoBiT assay. Luminescence data are expressed as % of receptor response (*see Materials and Methods*). Potency and efficacy of quinpirole dose-response curves in the absence or presence of NCS-1 were assessed. Results are expressed as mean ± SEM of four independent experiments each performed in triplicate. $^{**}p<0.01$, student's t test. **(B)** Table with summary of $pEC_{50}$, $\Delta pEC_{50}$, $E_{max}$ and $\Delta EC_{max}$ values derived from the NanoBiT assay. Source data are provided as a Source Data File.

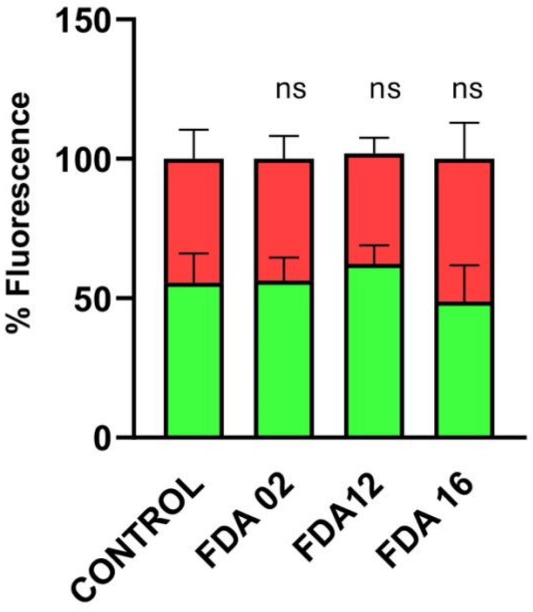


**Supplementary Figure 2:** **FDA Compounds have no effect in $D_2R$ cell localization.** Quantification of $D_2R$ fluorescence distribution between the membrane (red) and cytoplasm (green) in HEK293 cells transfected with $D_2R$ and treated with vehicle (Control) or 5 µM of the indicated FDA compounds. Statistical comparison was performed with Control. Bars represent the mean ± SD of five fields per condition from three independent experiments, ns: non-significant.

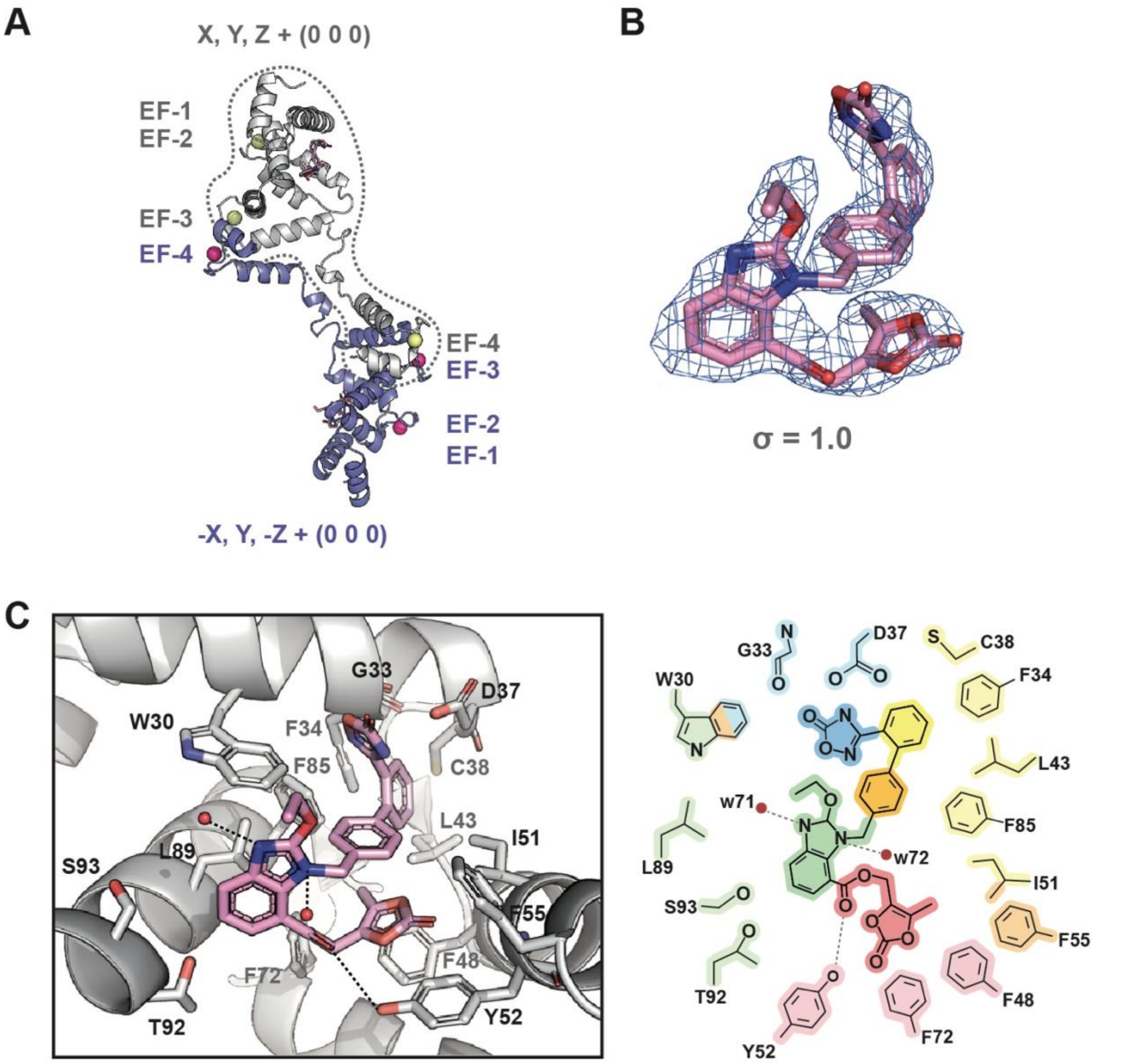


**Supplementary Figure 3:** Structural details on the complex of NCS-1 with azilsartan medoxomil **(A)** Representation of the domain swapping found. The NCS-1 molecule found in the A.U. is shown in grey and the symmetry related molecule with which the EF-4 motif is exchanged, in blue. **(B)** 2Fo-Fc electron density map. **(C)** Detail of the residues involved in AZS recognition. Hydrogen bonds between NCS-1 (grey) and AZS (pink) are represented as black dashed lines. Water molecules are shown as red spheres. Two-dimensional interaction diagram of the NCS-1/AZS F85complex. Residues are color-coded according to their interaction with different ligand moieties: oxadiazole (blue), phenyl rings (yellow and orange), benzimidazole (green), ester group and dioxolane (red).

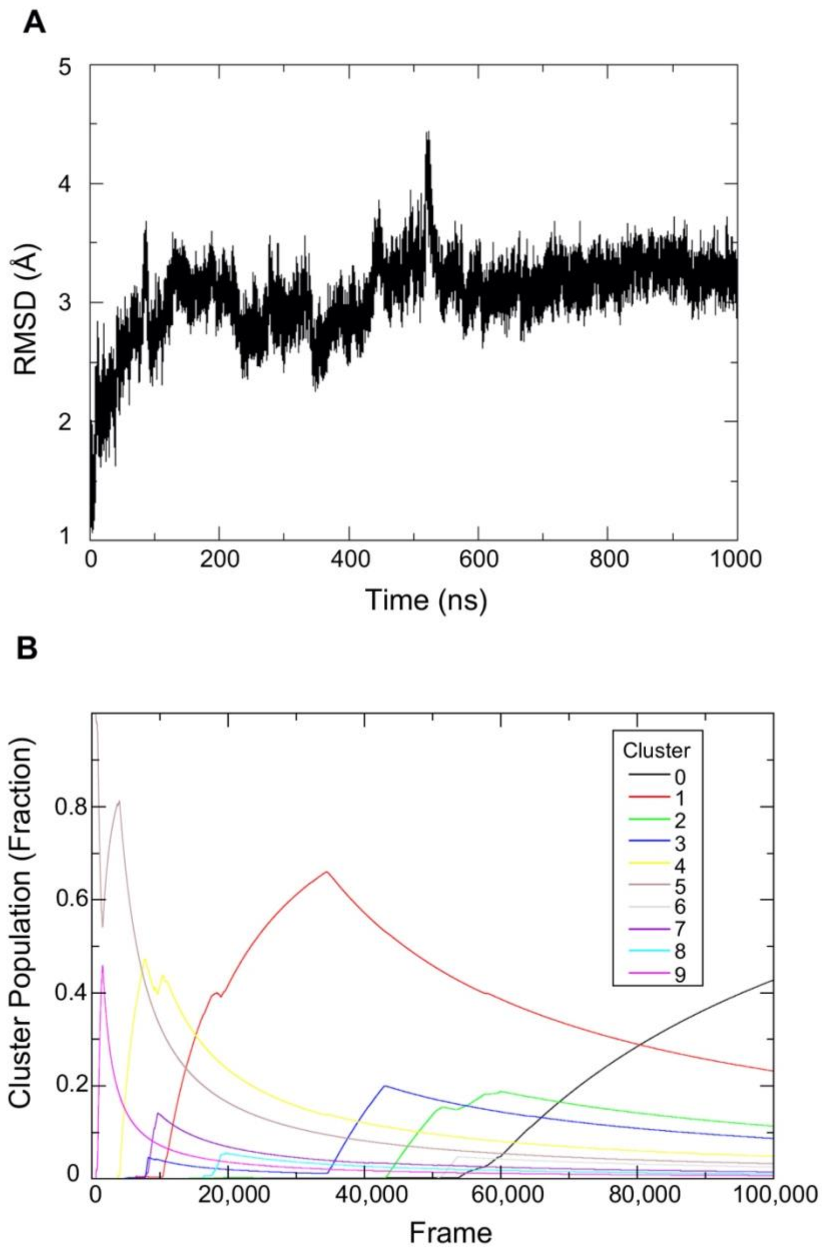


**Supplementary Figure 4:** Molecular dynamics simulations of the NCS-1/AZS complex. **(A)** Root-mean-square deviation (RMSD) analysis of the protein backbone atoms. **(B)** Hierarchical clustering of the MD trajectories using the average-linkage algorithm. The energy-minimized structure was used as the reference for aligning the MD trajectories prior to RMSD analysis.

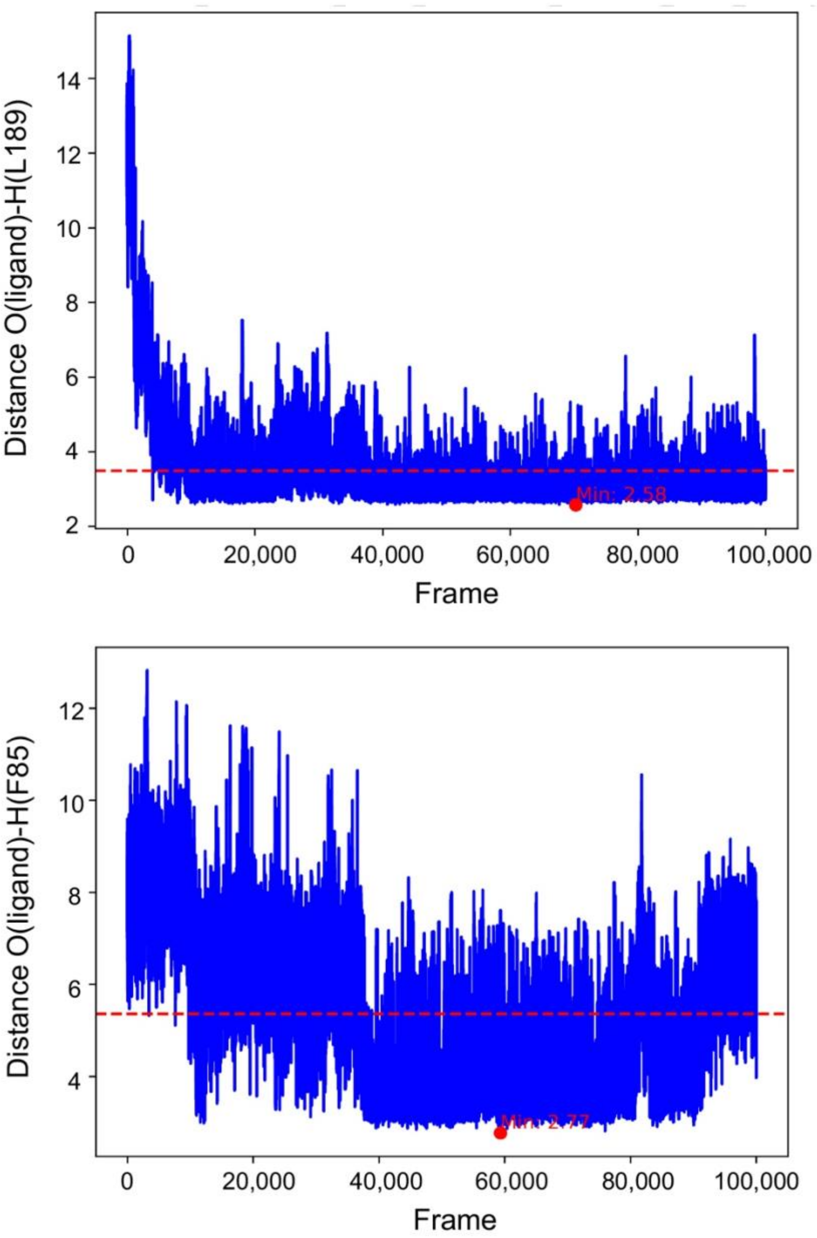


**Supplementary Figure 5:** Hydrogen bond distances between AZS and L189 or F85 along the molecular dynamics simulations. The mean value is represented by the dashed line, and the minimum value is also indicated.

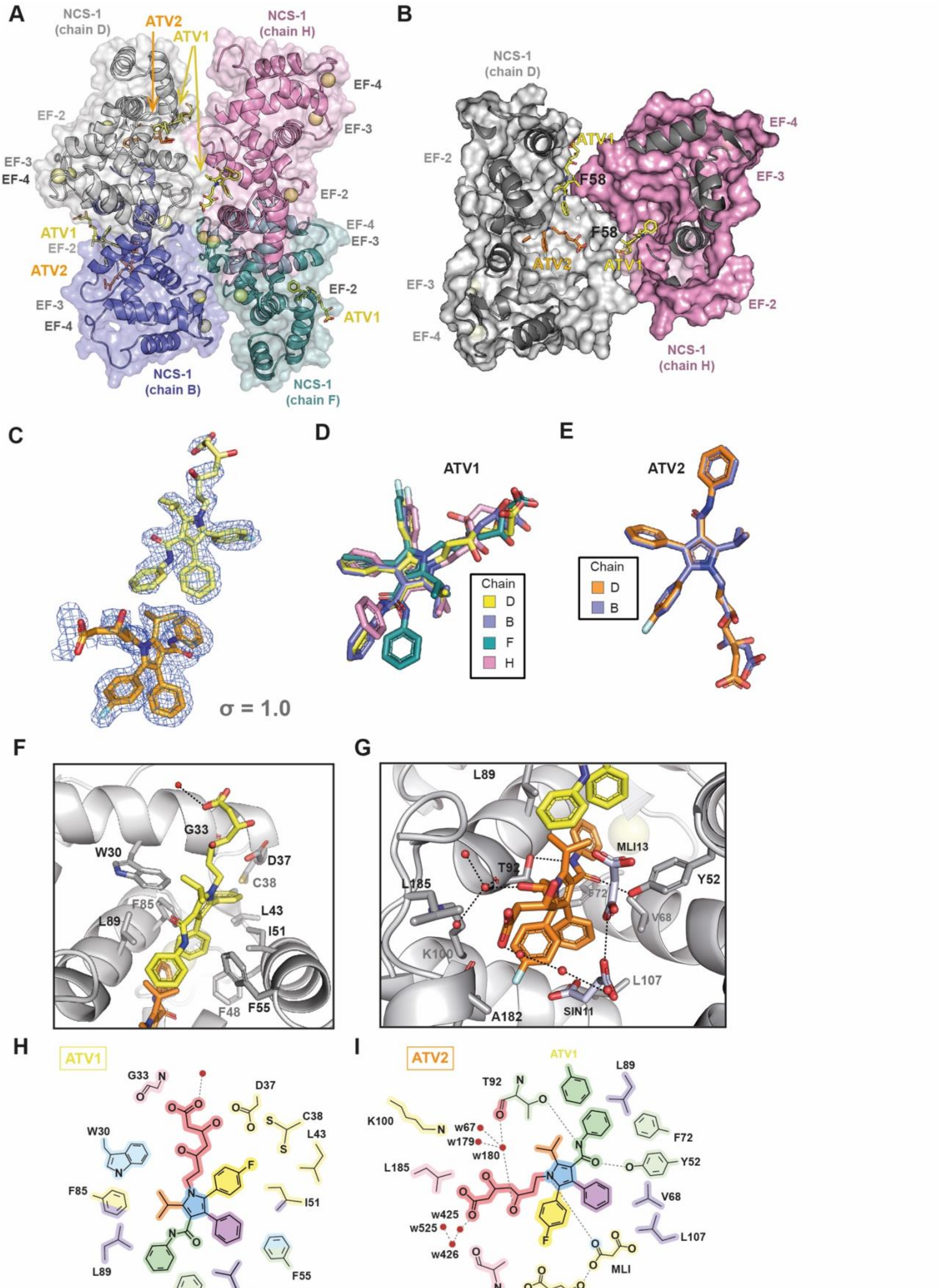
A
NCS-1 (chain D)
ATV2
ATV1
NCS-1 (chain H)
EF-4
EF-2
EF-3
EF-3
EF-4
EF-2
EF-4
EF-3
ATV1
EF-2
ATV2
EF-2
EF-3
ATV1
EF-4
NCS-1 (chain B)
NCS-1 (chain F)
B
NCS-1 (chain D)
ATV1
EF-4
EF-2
F58
EF-3
F58
ATV2
ATV1
EF-3
EF-2
EF-4
NCS-1 (chain H)
C
σ = 1.0
D
ATV1
Chain
D
B
F
H
E
ATV2
Chain
D
B
F
W30
G33
D37
C38
F85
L43
L89
I51
F48
F55
G
L89
MLI13
T92
L185
Y52
F72
V68
K100
L107
A182
SIN11
H
ATV1
G33
N
D37
C38
W30
L43
F85
F
I51
L89
F55
ATV2
F48
I
ATV2
ATV1
L89
T92
N
K100
N
w67
w179
w180
F72
Y52
L185
V68
w425
w525
w426
L107
MLI
A182
SIN

**Supplementary Figure 6:** Structural details of the complex of NCS-1 with atorvastatin **(A)-(E)** Analysis of the independent molecules comprising the asymmetric unit of the NCS-1/ATV crystal. **(A)** NCS-1 is shown as ribbons and colored according to chain ID: B (purple), D (gray), F (green), and H (pink). The ATV1 and ATV2 ligands are indicated and represented as sticks in yellow and orange, respectively. $Ca^{2+}$ ions are shown as yellow spheres. **(B)** Molecular surface and ribbon representation of NCS-1 chains D (gray) and H (pink). The positions of F58 residues, which act as lids over the ATV1 binding site, are indicated. **(C)** 2Fo-Fc electron density map corresponding to the two ligands (ATV1 and ATV2) bound to NCS-1 chain D, contoured at 1.0 σ. **(D)** and **(E)** Superposition of the ATV ligands bound to NCS-1 and modelled in the A.U. **(F)**, **(G)** Close-up of the residues involved in ATV recognition. Hydrogen bonds between NCS-1 (gray), ATV1 (yellow), and ATV2 (orange) are shown as black dashed lines. Water molecules are represented as red spheres. Malonate (MLI13) and succinate (SIN11) molecules involved in recognition are also depicted. **(H)**, **(I)** Two-dimensional schematic of the interactions between NCS-1 and ATV1 **(H)** and ATV2 **(I)**, respectively. Residues are color-coded according to their interaction with the different ligand groups: 3,5-dihydroxypentanoyl end (red), methylpropane (orange), fluorobenzene (yellow), phenyl (purple), benzamide (green), and pyrrole (blue).

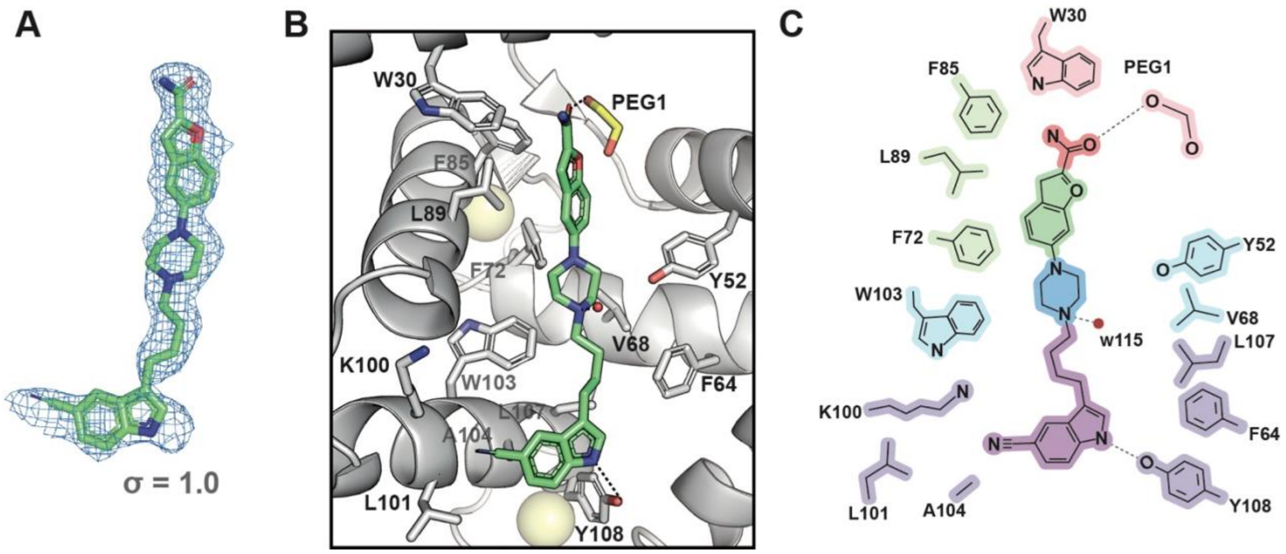


**Supplementary Figure 7:** Structural details of the complex between NCS-1 and vilazodone. **(A)** 2Fo-Fc electron density map corresponding to VLZ and contoured at 1.0 σ. **(B)** Interactions between NCS-1 (gray) and VLZ (green). The residues involved in ligand recognition are shown. Water molecules and $Ca^{2+}$ ions are represented as red and yellow spheres, respectively, and the PEG molecule is shown as yellow sticks. Hydrogen bonds between NCS-1 and VLZ are indicated with black dashed lines. **(C)** Two-dimensional interaction diagram. Residues are color-coded according to their interaction with the different ligand groups: amide (red), benzofuran (green), piperazine (blue), and the aliphatic chain, indole, and cyano groups (purple).

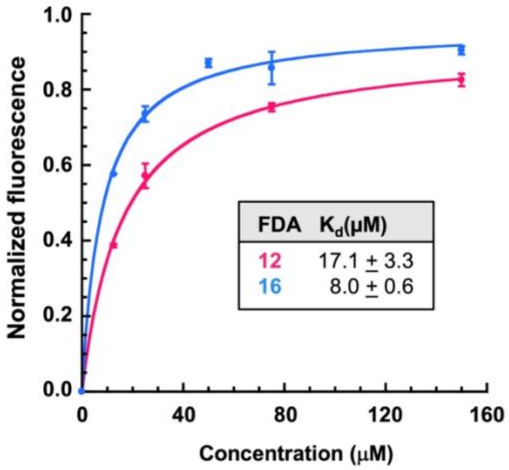


**Supplementary Figure 8: The binding of atorvastatin and vilazodone to NCS-1ΔH10:** Intrinsic emission fluorescence assays at increasing concentrations of the FDA compounds.

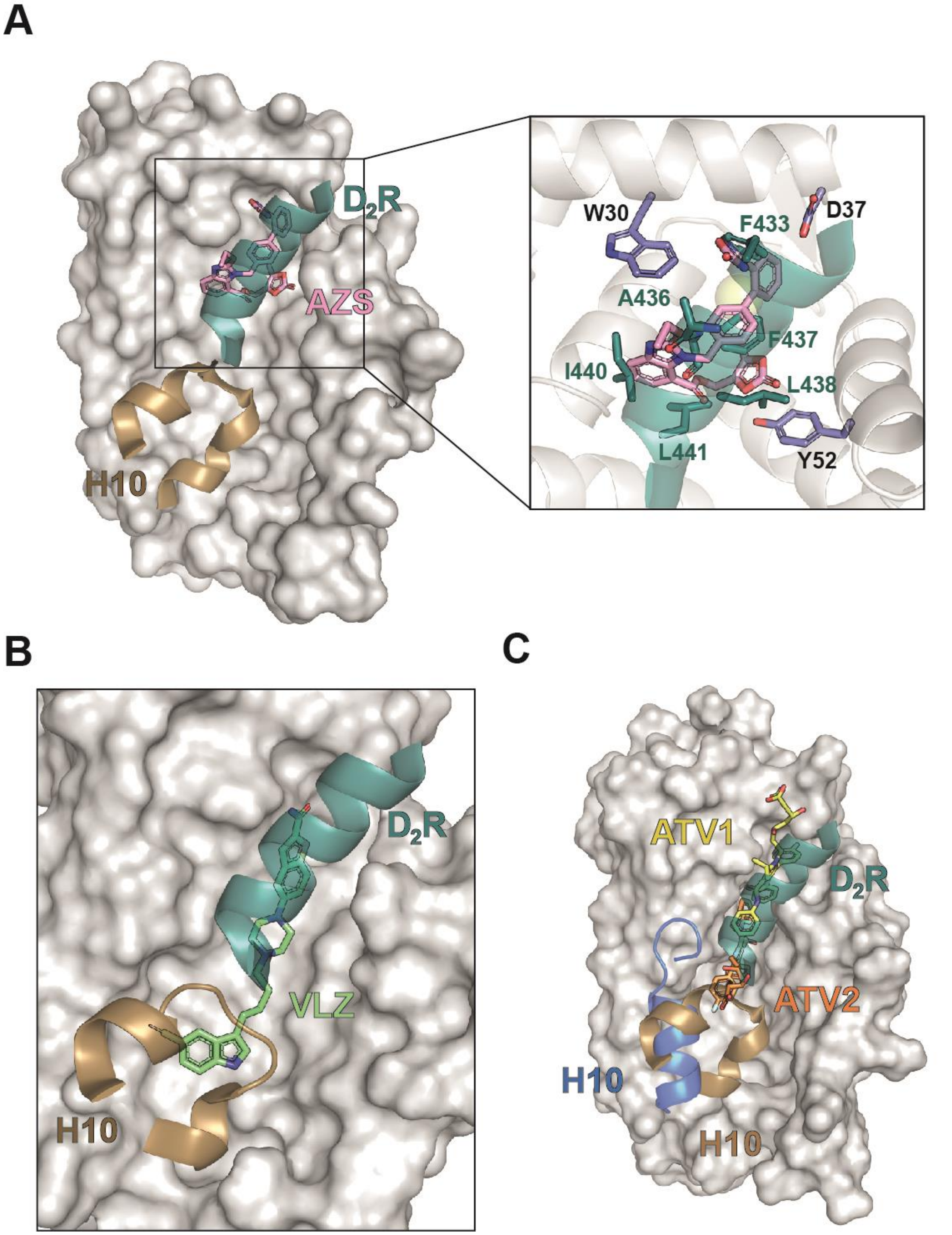


**Supplementary Figure 9: Structural comparison of NCS-1/FDA vs NCS-1/$D_2$R complexes:** The NCS-1/FDA complexes were superposed onto the NCS-1/$D_2$R H8 complex (PDB code: 5AER; (Pandalaneni et al., 2015)). The NCS-1/$D_2$R H8 complex is depicted as follows: NCS-1 as molecular surface or grey ribbons (with helix H10 highlighted in gold). Only The $D_2R$ helix H8 (green ribbon) that is placed in the upper part of the cavity is shown, since FDA drugs overlap only with this helix. In the case of the NCS-1/FDA structures, only the FDA compounds (sticks and color code as in Figure 8) and the NCS-1 helix H10 (if present, blue ribbon) are shown. **(A)** The NCS-1/AZS complex. A close-up view highlights how AZS positions its functional groups in regions where the $D_2R$ helix H8 places side chains (shown as green sticks) to mediate recognition by NCS-1 (interacting residues shown as lilac sticks). **(B)** and **(C)** The NCS-1/VLZ and NCS-1/ATV complexes, respectively.

**Supplementary Table 1: Top 20 FDA-approved compounds:** The original therapeutic indication, key residue interactions, docking scores, and MMGBSA binding energies are shown.

| Compound FDA-number | Target / Treatment / Crossing the blood-brain barrier | Interactions | QPlogPo/w | QPlogS |
|---|---|---|---|---|
| **Azilsartan** FDA-02 | Angiotensin II type 1 receptor antagonist / Antihypertensive / Low penetrance | 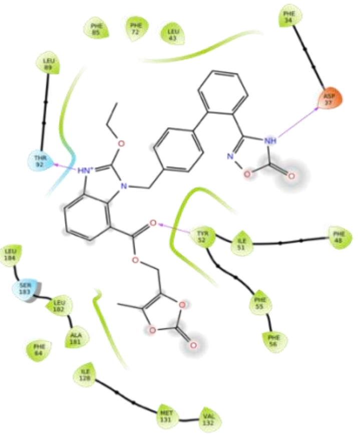 | 3.851 | -7.175 |
| **Sunitinib** FDA-03 | Multitargeted tyrosine kinase inhibitor (TKI) / Anticancer, kidney cancer / No | 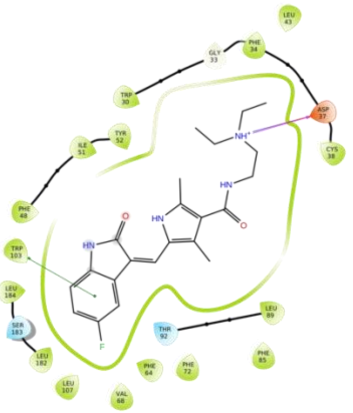 | 3.781 | -4.795 |

| | | | | |
|---|---|---|---|---|
| **Nelfinavir** **FDA-04** | HIV-1 protease inhibitor / HIV antiretroviral / No | 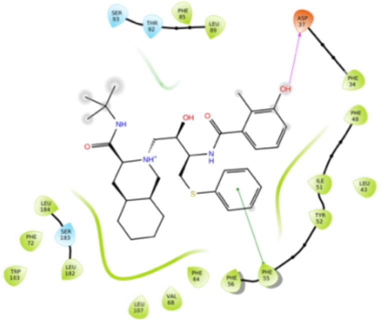 | 4.443 | -6.219 |
| **Ibutilide** **FDA-05** | Calcium channels / Antiarrhythmic / No | 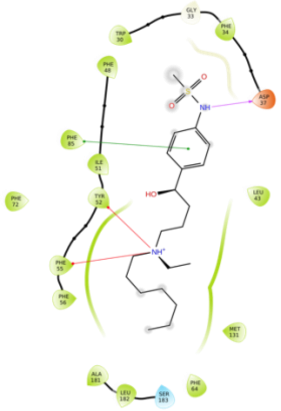 | 3.242 | -1.821 |
| **Lapatinib** **FDA-06** | Tyrosine kinase inhibitor / Anticancer, breast cancer / No | 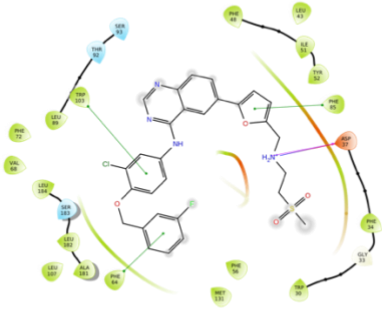 | 5.901 | -6.953 |

| | | | | |
|---|---|---|---|---|
| **Ergotamine** **FDA-08** | 5-HT adrenergic and serotonin receptor agonist / Vasoconstrictor, migraines / No | 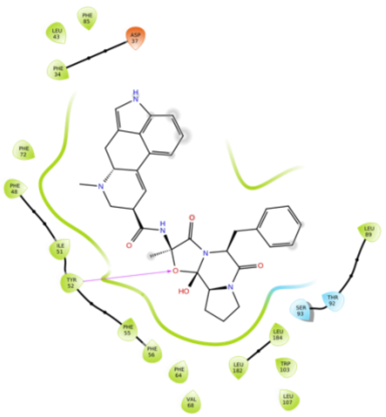 | 2.435 | -2.728 |
| **Darunavir** **FDA-09** | HIV-1 protease inhibitor / Antiretroviral for HIV / No | 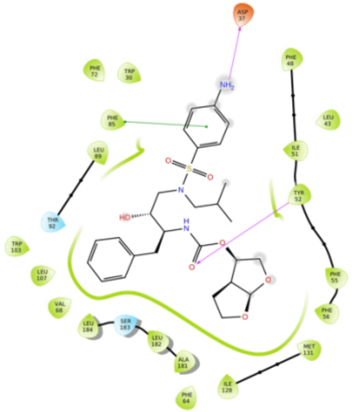 | 2.69 | -4.185 |

| | | | | |
|---|---|---|---|---|
| **Dabigatran** **FDA-10** | Prothrombin, liver carboxylesterase-1, cocaine esterase, UDP-glucuronosyltransferase / Anticoagulant / No | 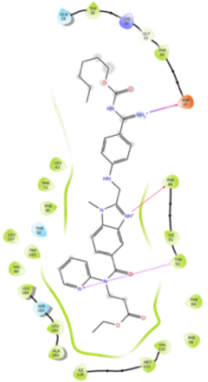 | 6.348 | -9.256 |
| **Faslodex** **FDA-11** | Estrogen receptor antagonist / Anticancer, breast cancer / No | 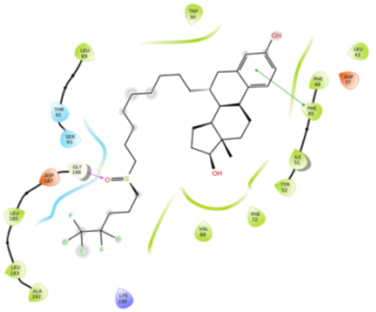 | 8.037 | -8.963 |
| **Atorvastatin** **FDA-12** | HMG-CoA reductase inhibitor / Dyslipidemia and cardiovascular disease prevention / Yes | 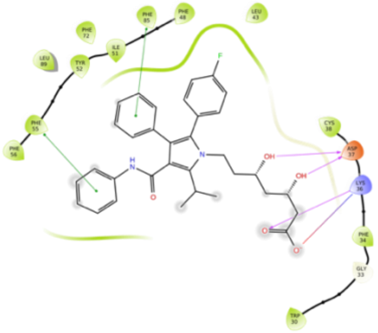 | 6.901 | -7.872 |

| | | | | |
|---|---|---|---|---|
| **Hydroxy-cloroquine**<br>**FDA-13** | RNA polymerase inhibitor; Toll-like receptors (TLRs) / Antimalarial and antirheumatic / Yes | 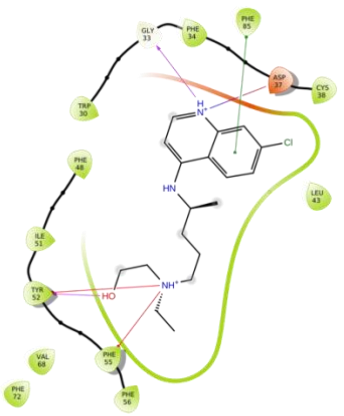 | 3.312 | -3.607 |
| **Cloroquine**<br>**FDA-14** | RNA polymerase inhibitor; Toll-like receptors (TLRs) / Antimalarial and antirheumatic / Yes | 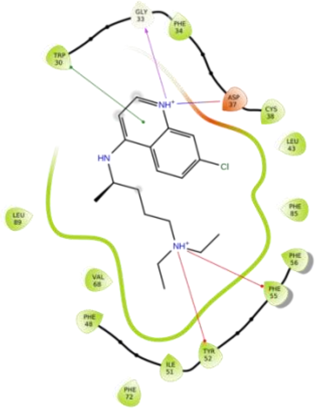 | 4.399 | -4.271 |

| | | | | |
|---|---|---|---|---|
| **Salmeterol** **FDA-15** | Ab2 adrenergic receptor agonist / Asthma /Yes | 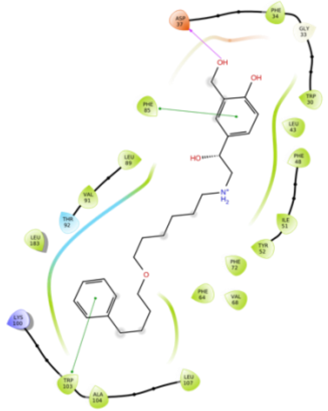 | 3.732 | -3.718 |
| **Vilazodone** **FDA-16** | Partial agonist of the serotonin 5-HT1A receptor and serotonin transporter inhibitor / Antidepressant / Yes | 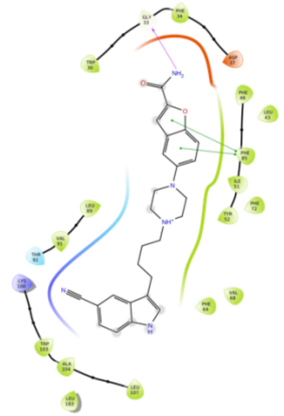 | 3.023 | -6.661 |

| | | | | |
|---|---|---|---|---|
| **Clomifene**<br>**FDA-17** | Estrogen receptor modulator / Female infertility / No | 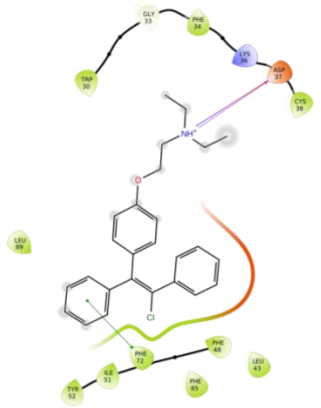 | 7.006 | -6.282 |
| **Ponatinib**<br>**FDA-18** | Third-generation multitargeted tyrosine kinase inhibitor / Chronic myeloid leukemia / Yes | 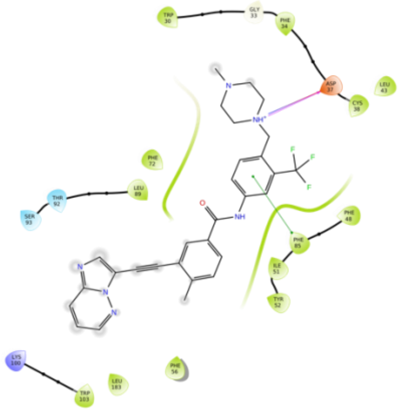 | 4.37 | -6.041 |

| | | | | |
|---|---|---|---|---|
| **Imatinib** FDA-19 | First-generation tyrosine kinase inhibitor / Chronic myeloid leukemia / Yes | 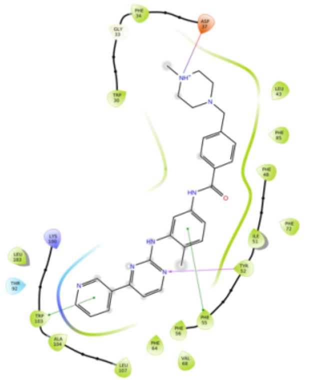 | 3.679 | -5.253 |
| **Daclatasvir** FDA-20 | Nonstructural protein 5A (NS5A) inhibitor / Antiviral, Hepatitis C virus / No | 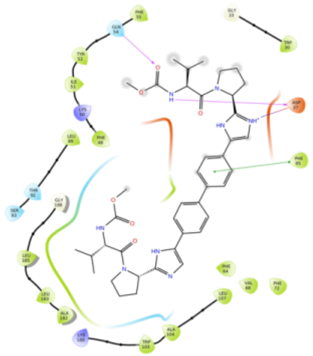 | 5.814 | -9.964 |
| **Protriptyline** FDA-22 | Acetylcholinesterase inhibitor / Antidepressant / Yes | 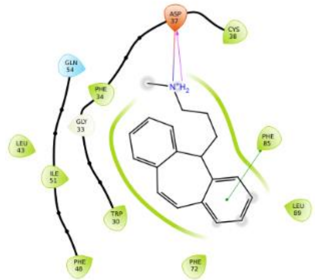 | 4.791 | -4.249 |

| | | | | |
|---|---|---|---|---|
| **Fluoxetine** **FDA-82** | Selective serotonin reuptake inhibitor / Antidepressant, bulimia, or bipolar disorder / Yes | 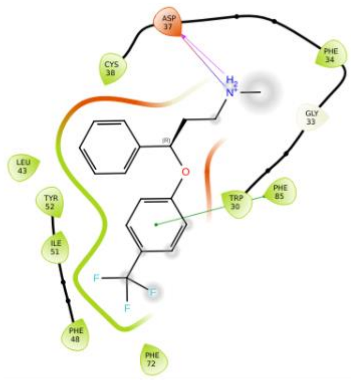 | 4.46 | -3.653 |
